# Still Not Sterile: Chlorhexidine gluconate treatment does not completely reduce skin microbial bioburden and promotes pathogen overabundance in patients undergoing elective surgeries

**DOI:** 10.1101/2024.07.20.602341

**Authors:** Elizabeth C. Townsend, Kayla Xu, Karinda De La Cruz, Lynda Huang, Shelby Sandstrom, Delanie Arend, Owen Gromek, John Scarborough, Anna Huttenlocher, Angela L.F. Gibson, Lindsay R. Kalan

**Author notes:** Lindsay Kalan 1280 Main St W HSC-4H45 Hamilton, ON, L8S 4L8.

## Abstract

Surgical site infections (SSI) continue to occur despite widespread adoption of surgical antiseptics. The effects of chlorhexidine gluconate (CHG)-based antiseptics on the skin microbiome also remains undefined due to confounding effects of CHG persistence on skin. Patients undergoing elective surgery were enrolled to characterize the immediate and long-term impact of pre-surgical preparation with CHG antiseptic on skin microbial communities. Due to the broad-spectrum antimicrobial activity of CHG and its propensity to bind extracellular DNA, methods to selectively identify live microorganisms are critical to this process and to fully elucidate the effectiveness of pre-surgical protocols and potential disruptions to the healthy skin microbiome. Swabs of the surgical site skin microbiome were collected at multiple timepoints before and after surgery. Microbial bioburden and community compositions were evaluated with viability qPCR and 16S ribosomal RNA gene profiling. Pre-operative CHG induced a measurable reduction in the viable microbial bioburden at the surgical site. On the day of surgery, surgical sites displayed a significant increase in the relative abundance of several SSI associated bacterial genera, including*, Acinetobacter, Bacillus, Escherichia-Shigella,* and *Pseudomonas*, compared to baseline*. Bacillus* species isolated from subjects at baseline showed resistance to CHG with MICs exceeding 1000 µg/ml. Despite major shifts in the skin microbiome upon exposure to CHG, they were transient in the majority of individuals. Skin microbial community structure recovered by the post-surgical follow-up. In short, this study shows that pre-surgical application of CHG can significantly reduce viable skin microbial bioburden, however, complete sterility is not achieved. While CHG induces temporary shifts in the skin microbiome, including enrichment for potentially pathogenic taxa, the skin microbiome recovers back to near baseline. Collectively, these findings identify tangible avenues for improving antiseptic formulations and offer further support that the skin microbiome is viable, stable, and resilient to chemical perturbation.

## Introduction

Surgical site infections (SSI) pose a substantial burden to affected patients and the healthcare system. Despite widespread adoption of broad-spectrum antiseptics to reduce the skin microbial bioburden at the time of surgery, SSI still occur. Approximately 0.5% to 3% of all surgical patients will experience an infection at or adjacent to their surgical incision.^1,2^ Collectively, these infections pose an additional $3.5-$10 billion in annual healthcare costs due to prolonged hospital stays, additional diagnostic tests, procedures and operations, and increased utilization of outpatient resources.^3,4^ To prevent SSI, antiseptics intentionally reduce skin microbial bioburden at the time of surgery, but in the process disrupt the skin’s naturally occurring microbial communities. Defining the impact of these antiseptics on the skin microbiome across various surgeries and surgical sites is needed.

The skin microbiome comprises complex microecosystems of bacteria, fungi, and viruses.^5–7^ The density of eccrine, apocrine, and sebaceous glands across the body create moist, sebaceous, or dry microenvironments, which respectively promote site-specific microbial populations based on the local physiology. Under normal healthy conditions, the skin microbiome promotes skin health by enhancing barrier function and preventing pathogen overgrowth through competitive interactions and niche exclusion.^5,7,8^ There is currently a major gap in our understanding of the differential impacts of antiseptics on skin microbial communities at healthy and surgical sites, including potential selective pressures promoting antiseptic resistance. Perhaps most important is to understand how healthy skin communities recover following the major chemical perturbation that occurs after antiseptic application.

Standard pre-surgical preparations often involve having patients shower or bathe with 4% chlorhexidine gluconate (CHG) soap the night before and morning of surgery, as well as locally applying CHG (e.g. ChloraPrep) to the surgical site just before incision.^9–12^ CHG and other topical antiseptics are validated for clinical use through demonstrating a significant reduction in culturable microbial bioburden for up to 48 hours post-application.^13–20^ However, the impact of broad spectrum antiseptics like CHG, on healthy skin microbial communities remains unknown, despite its widespread clinical use. Previous sequencing-based efforts to quantify and characterize the impact of CHG on the skin microbial communities have yielded mixed results. While some demonstrate a reduction in microbial diversity following CHG exposure,^21^ most studies conclude that no significant change in skin microbial community structure or bioburden occurs.^22–25^ It has been proposed that this discrepancy between culture-dependent and -independent studies is due to CHG’s ability to bind persistent bacterial DNA to the surface of the skin.^22,26^ CHG is a highly positively-charged molecule that disrupts negatively-charged microbial cell membranes.^26^ This property likely allows CHG to bind DNA and proteins released from the recently lysed cells and upper skin layers,^22,26^ which subsequently confounds sequencing based analyses that do not account for the viability of sampled microbes. This underscores the pressing need for establishing a method capable of accurately evaluating live microorganisms on the skin, particularly after a toxic (eg. antiseptic) exposure, via DNA sequencing-based technologies.

With this work, we aim to characterize the immediate and long-term impact of CHG antiseptic on viable skin microbial bioburden and community composition in patients undergoing elective surgery. To selectively evaluate DNA from live microorganisms within the skin microbiome pre- and post-antiseptic exposure, we optimized a propidium monoazide (PMAxx) based viability assay. Previous work has used PMAxx to estimate the viable portion of the skin microbiome,^27^ concluding that viable bacterial cells are very low. Here, we show that the concentrations of PMAxx previously reported are toxic towards live bacteria. We determined the optimal concentration of PMAxx for evaluating viable skin bacterial communities. We show that most bacterial cells collected from skin microbiome samples are viable and pre-operative preparation with CHG effectively reduces viable microbial bioburden at the surgical site. Concurrently, we see enrichment of potentially pathogenic taxa. We further demonstrate that these shifts are transient, with the majority of individuals’ skin microbiomes recovering, both in absolute abundance and microbial community structure, by their post-surgical clinic follow-up. *Bacillus* and *Enterococcus* species isolated from the skin of these surgical patients also display increased resistance to CHG. Collectively these findings identify avenues to spur the development of improved pre-surgical protocols and broaden our understanding of skin microbiome dynamics.

## Methods

### Subject Identification and Enrollment

Adults 18 years or older undergoing elective surgery were enrolled from the UW-Health General Surgery Clinic. Inclusion criteria for the study; i) person 18 years or older, ii) undergoing elective surgery requiring preoperative CHG, iii) surgical wound anticipated to be classified as clean, clean-contaminated, or contaminated, and iv) able to provide written and verbal consent. Exclusion criteria: i) emergent surgery or surgery due to trauma, ii) recent burn encompassing more than 30% of total body surface area, iii) surgical wound anticipated to be classified as dirty-infected, iv) known allergy to chlorhexidine or v) administered antibiotics (not including those for surgical prophylaxis) within 2 weeks of enrollment or the surgery. Information related to pre-surgical assessments, surgery, post-surgical care, following post-operative outpatient evaluations and patient co-morbidities were extracted from the medical record. Significant alcohol use was defined as a recorded history of alcoholism or consuming 14 or more drinks per week at the time of enrollment. If a subject developed a surgical site infection or other surgical site occurrence developed, information related to its assessment and care were also recorded.

### Survey of Subject Home and Work Environment

To gauge environmental factors that may influence subject skin microbial communities, subjects completed a survey to collect information on their skin health, hygiene habits, and home and work environments. This survey was administered by study staff at the initial clinic visit following enrollment. Questions pertained to: i) select skin and medical history that may not be apparent in their medical chart (e.g. do they consider their skin to generally be oily, dry, or a combination), ii) personal hygiene habits (e.g. number of showers per week), iii) home environment (e.g. do they live in a urban, suburban or rural setting), iv) work environment (e.g. do they work in a health care setting), and v) exposure to animals (e.g. do they live with cats, dogs, or other animals).

### Pre-surgical Preparation

Leading up to surgery, subjects followed the UW-Health guidelines for pre-surgical preparation. Subjects were provided 4% CHG soap (Hibiclens; Mölnlycke Health Care, Norcross, Georgia) to take home and shower with the night prior and morning of their surgery following the instructions outlined in the UW-Health pre-surgical guidelines.^9^ When patients arrived into the pre-operative waiting area, patients self-reported number of showers / baths patients completed with CHG prior to their surgery and whether they used other hygiene products (e.g. shampoos, body wash) for these showers. In the operating room, application of a CHG based antiseptic occurred according to surgical guidelines. In general, a 26ML applicator of 2% CHG with 70% Isopropanol (ChloraPrep; Becton, Dickinson and Company (BD), Franklin Lakes, NJ) was gently pressed to the surgical field until the solution was visible and applied with repeated up and down and back and forth strokes or a circular motion extending outwards for at least 30 seconds. The umbilicus was cleaned with a swab of CHG if applicable. CHG solution was left to air dry for at least 3 minutes prior to the sample collection and subsequent incision.

### Sample Collection

Swabs of the skin microbiome were collected from the intended surgical site (pre-surgery) / around the incision (after surgery) and a similar control site that was not anticipated to receive CHG application in the operating room. The armpit served as the control site for surgeries at moist body sites (e.g. inguinal or umbilical hernia repairs), and the volar forearm served as the control site for surgeries at dry body sites (e.g. cholecystectomies or ventral hernia repairs). If CHG or another antiseptic had been applied to the volar forearm for IV insertion, the contralateral forearm was used. **Figure 1A** illustrates example sampling locations.

**Figure 1:**
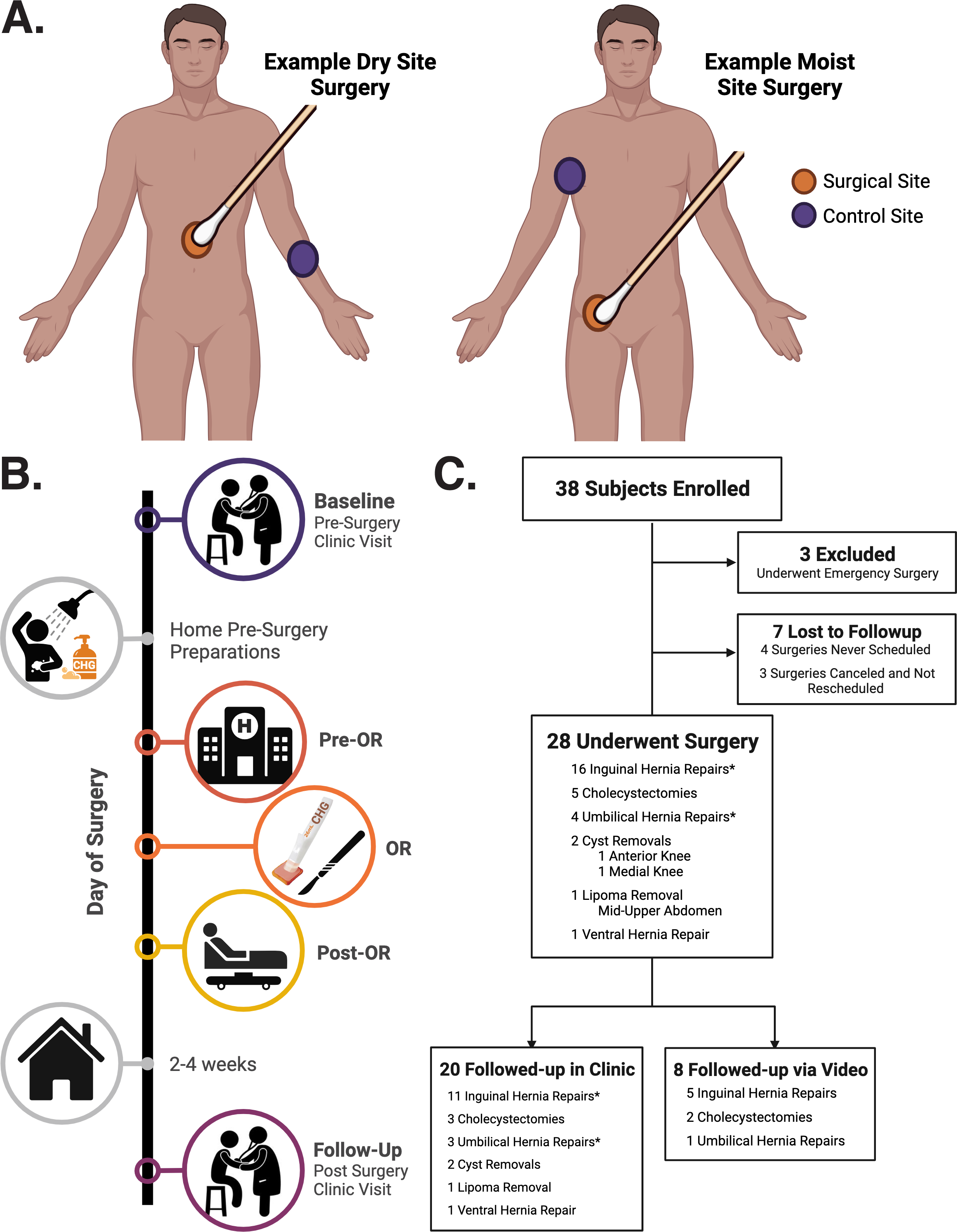
Study Design. A. Skin microbiome sampling locations for dry skin site surgeries (e.g. central or upper abdomen, knee; left image) and moist skin site surgeries (e.g. lower abdomen to inguinal crease or umbilicus; right image). **B.** Sample collection timeline. **C.** Flow chart illustrating the number of enrolled subjects that were excluded or underwent elective surgery and had their post-surgery clinic visit in person of over video. * Indicates one subject underwent simultaneous inguinal and umbilical hernia repair. For this subject, both surgery sites were sampled at each timepoint.

Swabs were collected from both the surgical site and a control site at multiple timepoints (**Fig. 1B**). These include; i) at the pre-surgery evaluation clinic visit, ii) in the pre-operative care unit, in the operating room after CHG was applied and allowed to dry for 3 minutes yet prior to surgical incision, and in the post-anesthesia care unit before discharge, and iii) at the post-surgery clinic visit. Swabs designated for DNA extraction were placed into 155 μl of 1% Bovine Serum Albumin (BSA) in 1x Phosphate Buffered Saline (PBS) and were stored at 4°C for less than 2 hours before processing for selective detection of DNA from viable organisms (per below) and eventual DNA extraction. Swabs designated for microbial culture were placed into 1 ml of liquid Amies (Copan Diagnostics Inc., Murrieta, CA). Culture swabs were stored at 4°C for less than 2 hours before being processed for microbial culture.

### Microbial Culture and Bacterial Isolate Identification

Swabs designated for microbial culture were spun down using DNA IQ Spin Baskets (Promega, Madison, WI). A portion of each sample was serially diluted with 1X phosphate buffered saline (PBS) and plated onto Tryptic Soy Agar (TSA) with 5% sheep blood (BBL, Sparks, MD), Brain Heart Infusion (BHI) agar, and BHI agar with 1% Tween 80 and 50mg/L mupirocin for quantitative bacterial culture. Plates were incubated at 35°C overnight. To isolate culturable bacteria, colonies with distinct morphology were isolated and incubated at 35°C overnight on the same media as the original culture or BHI with tween without mupirocin, then single colonies were inoculated into liquid Tryptic Soy Broth (TSB), BHI broth, or BHI broth with 1% tween for overnight incubation. To identify each bacterial isolate, a portion of the overnight culture underwent DNA extraction and sanger sequencing (Functional Biosciences, Madison, WI) of the bacterial 16S ribosomal RNA gene. The remaining portion of the isolate culture was stored in glycerol at -80°C.

### Measuring CHG Resistance in Bacterial Isolates

Bacterial isolates collected from surgical subjects during the baseline clinic visit were evaluated for resistance to CHG via a disc diffusion minimum inhibitory concentration (MIC) assay. Isolates were grown overnight in BHI with 1% Tween 80 liquid media at 37°C. Overnight cultures were diluted with 1X PBS using optical density readings to achieve 1x10^8^ bacterial colony forming units / ml. Diluted cultures were plated to create a bacterial lawn on BHI agar with 1% Tween 80 and plates were dried for 5-10 minutes prior to the placement of 6mm Whatman Antibiotic Assay Disc (Cytivia, Buckinghamshire, United Kingdom). 2% CHG in 70% Isopropanol (ChloraPrep; BD) was serially diluted with 1X PBS, 15 µl of each dilution was added to a respective disc to achieve a concentration range of 1-2000 µg/ml CHG. A disc with 1xPBS served as the negative control. Plates were then incubated for 18 hours overnight at 37°C. For each isolate, the disc with the smallest zone of inhibition, either just under the disc or extending no more than 1-3 millimeters beyond the disc was considered the MIC of CHG. All isolates were tested in duplicate and the average of the replicate MICs was calculated.

### Selective Detection of DNA from Viable Microorganisms

Swabs collected into 1% BSA in PBS were spun down using DNA IQ Spin Baskets (Promega, Madison, WI) and each sample was split into two equal 75 µl portions. To quantify and sequence the DNA from all (live and dead) members of the skin microbial communities, one portion of each sample was placed directly into -20C storage. To selectively quantify and sequence DNA from only live microbes within skin microbial communities, the other portion of each sample was processed with a modified propidium monoazide (PMAxx, Biotium, Fremont, CA) based viability assay (**Fig. 2A**). All steps involving PMAxx were done in a dark room. PMAxx was added to achieve a final concentration of 10 µM in each sample. Samples were rocked at room temp for 10 minutes, exposed to blue light for 15 minutes via the PMA-Lite LED Photolysis Device (Biotium, Fremont, CA), then spun at 5000 g for 10 minutes. Both the PMAxx treated portion and untreated portion of each sample were stored at -20 before DNA extraction.

**Figure 2:**
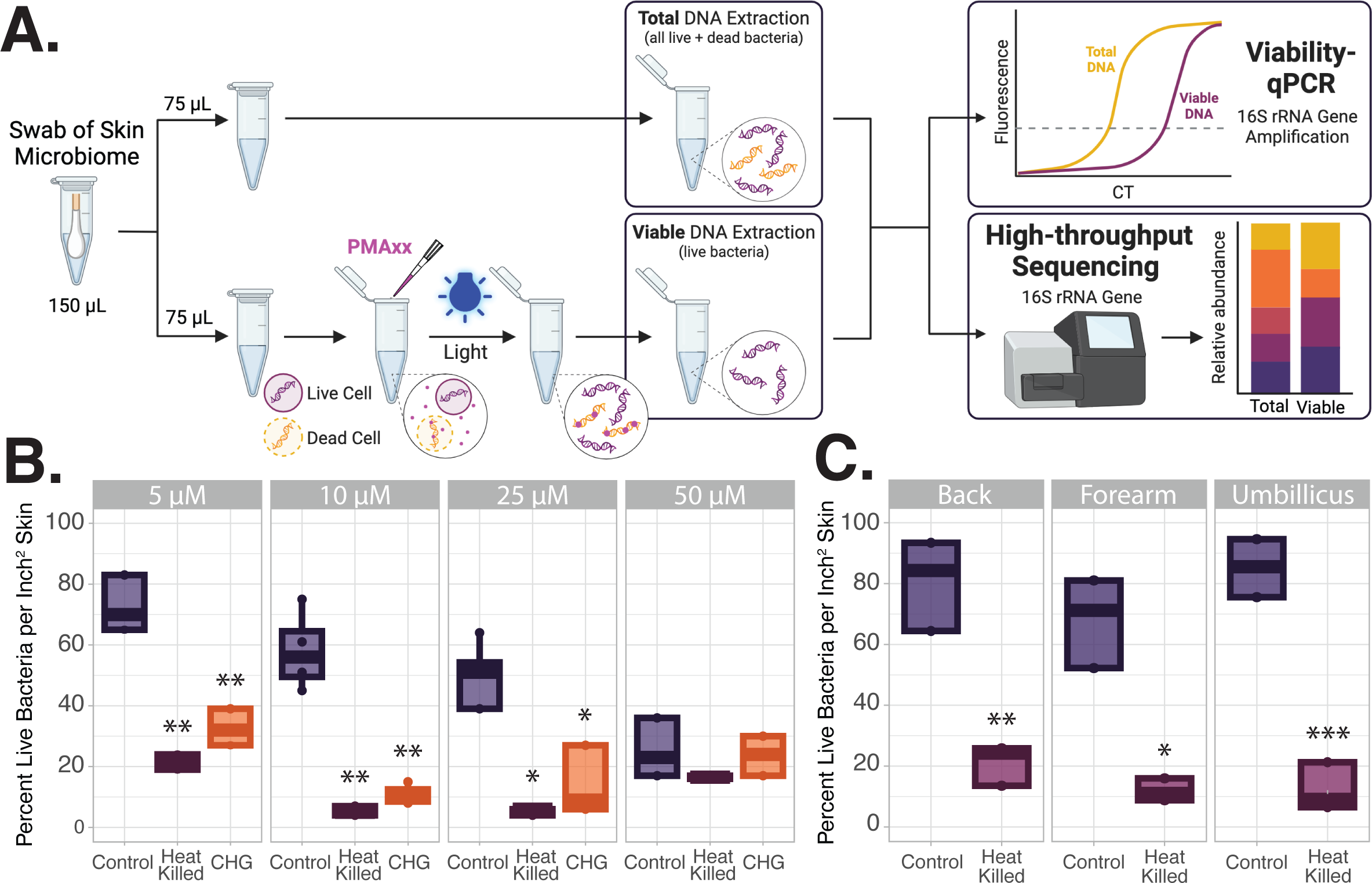
10 µM is an optimal PMAxx concentration for selective evaluation of live bacteria within skin microbial communities. A. Diagram of sample processing. Skin microbiome samples were split. Half of each sample was treated with PMAxx, which irreversibly binds to free DNA or DNA within compromised cells, allowing for selective amplification of only DNA from viable bacteria. The remaining half of the sample remained untreated to evaluate the DNA from both live and dead bacteria (Total DNA). Total and viable bioburden was evaluated via quantitative PCR of the bacterial 16S Ribosomal RNA gene (viability – qPCR). Viable and total microbial communities were further evaluated via high-throughput sequencing. **B.** Four different concentrations of PMAxx were evaluated for accurate quantification of viable and total bacterial bioburden on the volar forearm. Swabs of the forearm microbiome were collected from volunteers; one sample was heat killed at 95C for 10 minutes and one sample was collected following application of CHG (n=4 for 10µM, n=3 for remaining concentrations). Bioburden was quantified via viability-qPCR. Percentage of live bacteria was calculated by dividing the number of viable bacteria in the sample by the total number of bacteria. The percent live bacteria in heat killed samples and samples from CHG treated skin were compared to the control group by t-test with Welch’s correction. **C.** 10 µM of PMAxx accurately distinguishes control and heat killed microbial communities from multiple skin sites. Plot displays the percent live bacteria in a control and heat killed microbiome samples from the back, forearm, and umbilicus (n=3). Groups compared with via t-test with welches correction. * p-value < 0.05 ** p-value < 0.01 *** p-value < 0.001 **** p-value < 0.0001

Initial experiments aimed to optimize the viability assay parameters for selective evaluation of viable microbes within the complex microbial communities residing on skin both in control conditions and following application of CHG antiseptic. Swabs of the skin microbiome were collected from the volar forearm of healthy volunteers. Two samples were taken from one arm, one served as a control to determine the anticipated amount of viable bacteria on the skin in normal circumstances, and the other sample was boiled at 95°C for 10 minutes to heat-kill the majority of the bacteria present, as a negative control. The other forearm was treated with 2% CHG in 70% isopropanol prior to sample collection. Bacterial bioburden determined by qPCR of the 16S rRNA gene. The number of bacteria in the viable sample portion was divided by the number of bacteria in the total (live + dead) sample portion to determine the percentage of live bacteria within the sample.

### DNA Extraction, Library Construction, and Sequencing

DNA extraction was performed as previously described with minor modifications.^28^ Briefly, 300 μl of yeast cell lysis solution (from Epicentre MasterPure Yeast DNA Purification kit), 0.3 μl of 31,500 U/μl ReadyLyse Lysozyme solution (Epicentre, Lucigen, Middleton, WI), 5 μl of 1 mg/ml mutanolysin (M9901, Sigma-Aldrich, St. Louis, MO), and 1.5 μl of 5 mg/ml lysostaphin (L7386, Sigma-Aldrich, St. Louis, MO) was added to 150 μl of swab liquid before incubation for one hour at 37°C with shaking. Samples were transferred to a 2 ml tube with 0.5 mm glass beads (Qiagen, Germantown, Maryland) and bead beat for 10 min at maximum speed on a Vortex-Genie 2 (Scientific Industries, Bohemia, NY), followed by a 30 min incubation at 65°C with shaking, 5 min incubation on ice. The sample was spun down at 10,000 rcf for 1 min and the supernatant was added to 150 μl of protein precipitation reagent (Epicentre, Lucigen, Middleton, WI) and vortexed for 10s. Samples were spun down at maximum speed (∼21,000 rcf) and allowed to incubate at RT for 5 min. The resulting supernatant was mixed with 500 μl isopropanol and applied to a column from the PureLink Genomic DNA Mini Kit (Invitrogen, Waltham, MA) for DNA purification using the recommended protocol.

Viability quantitative polymerase chain reaction (viability-qPCR) was performed, to determine the amount of DNA from viable bacteria (treated with PMAxx) and total DNA from both live and dead bacteria (non-PMAxx treated) portion of each sample. In short, 1 µl of extracted DNA was added to a reaction mix containing 5 µl TaqMan Fast Advanced 2X Master Mix (Applied Biosystems, Waltham, MA), 0.5 µl TaqPman 16S 20X Gene Expression Assay with FAM (Applied Biosystems), and 3.5 µl PCR pure water. Samples were run for 40 thermos-cycles on the QuantStudio 7 Flex Real-Time PCR System (Applied Biosystems). Sample DNA concentrations were determined based on a standard curve of 0.015 to 15000 pg/µl DNA extracted from *Escherichia coli* (ATCC 1496).

16S rRNA gene V3-V4 region amplicon libraries were constructed using a dual-indexing method at the University of Wisconsin Biotechnology Center and sequenced on a MiSeq with a 2x300 bp run format (Illumina, San Diego, CA). Reagent-only negative controls were carried through the DNA extraction and sequencing process. A 20-Strain Staggered Mix Genomic Material (ATCC, Manassas, VA) served as a positive sequencing control.

### Sequence Analysis

The QIIME2^29^ environment was used to process DNA-based 16S rRNA gene amplicon data. Paired end reads were trimmed, quality filtered, and merged into amplicon sequence variants (ASVs) using DADA2. Taxonomy was assigned to ASVs using a naive Bayes classifier pre-trained on full length 16S rRNA gene 99% OTU reference sequences from the SILVA SSU database (release 138). Using the qiime2R package, data was imported into RStudio (version 1.4.1106) running R (version 4.1.0) for further analysis using the phyloseq package.^30^ Negative DNA extraction and sequencing controls were evaluated and removed from all samples based on absolute read count and ASV distribution in true patient samples. Abundances were normalized proportionally to total reads per sample. Data was imported into RStudio running R (version 4.2.1) for analysis. Plots were produced using the ggplot2 package. Taxa below 0.5% relative abundance were pooled into an “Other” category for the relative abundance plots. The Shannon alpha diversity index was used to measure the microbial diversity within individual samples. Bray-Curtis beta diversity metric was utilized to compare sample microbial community structures and all associated plots were ordinated via Non-metric Multidimensional Scaling (NMDS). Univariate and/or multivariate permutational multivariate analysis of variance (PERMANOVA) were used to evaluate associations between microbial community compositions and subject features. Each PERMANOVA was run considering the marginal effects of terms with 9999 permutations using Adonis2 in the vegan r package.^31^ ASVs from the *Staphylococcus* genus were aligned to the BLAST nucleotide database (2.14.1+) to assign probable species. MAASLIN2^32^ was utilized to identify significant differences in taxa abundance between various groups.

### Statistical Analyses

Most statistical analyses were conducted in R studio running R (version 4.2.1). Comparisons of subject demographics and co-morbidities between patient groups were analyzed via Prism (version 9.2.0).

## Data Availability

Sequence reads for this project can be found under NCBI BioProject PRJNA1092813. Code for analysis and generation of figures can be found on GitHub at https://github.com/Kalan-Lab/Townsend_etal_StillNotSterile

## Results

### Subjects

Thirty-eight subjects anticipated to undergo elective surgeries were enrolled (**Fig. 1C**, **Table 1**). Ten individuals were excluded because the surgery was never scheduled, the surgery was canceled and not rescheduled, or they underwent emergency surgery. The remaining individuals ranged in age from 30 to 81 years old (mean 58 ± 15 years). Of the subjects who underwent surgery, 19 had surgeries at moist body sites, these surgeries included inguinal hernia repairs (lower abdomen to inguinal crease) and umbilical hernia repairs. Nine subjects had surgeries at dry body sites, which included cholecystectomies, ventral abdominal hernia repairs, lipoma removal from the abdomen, and cyst removal from the knee. The proportion of subjects with co-morbidities and self-reported home and work environmental conditions were not significantly different between the two surgical sub-groups (**Table S1**). The surgical wound classification was considered clean for 21 (75%) subjects, clean-contaminated for 6 (21%; all the cholecystectomies and one umbilical hernia repair), and contaminated-infected for 1 subject (umbilical hernia repair). Twenty-four of the 28 subjects (85.7%) received antibiotic prophylaxis with a single dose of intravenous cefazolin the day of surgery (**Table S1**). Compared to the dry site surgery sub-group, a significantly higher proportion of those in the moist site surgery sub-group were male (90% vs. 44%, p-value < 0.05, Fishers exact), had lower BMIs (27.6 ± 3.4 vs. 33.9 ± 11.2, p-value < 0.05, Mann-Whitney), and had mesh implanted during the surgery (84% vs. 0%, p-value < 0.001, Fishers exact; **Table 1, Table S1**). Twenty of the subjects came to clinic for their follow-up 2-4 weeks post-surgery, while 8 conducted their follow-up over video and the final “Follow-up” skin microbiome samples were subsequently unable to be collected (**Fig. 1C**).

**Table 1:** Subject demographics; including average age, gender, self-reported race and ethnicity for the 38 enrolled subjects. Demographics are also broken down for the 10 subjects who were excluded of lost to follow-up, the 28 subjects who ultimately underwent surgery collectively as well as the 9 and 19 subjects, respectively, who underwent surgeries at dry or moist body sites. ∘ p-value < 0.05 for subjects that underwent surgery vs. excluded, fishers exact t-test. * p-value < 0.05 for subjects who underwent moist skin site surgeries vs. dry site surgeries, fishers exact t-test.

|  | All Subjects | Excluded or Lost to Follow-up | All Subjects who Underwent Surgery | Dry Site Surgeries | Moist Site Surgeries |
| --- | --- | --- | --- | --- | --- |
| **Basic Demographics** |  |  |  |  |  |
| Number subjects | 38 | 10 | 28 | 9 | 19 |
| Age ± SD | 57.3 ± 15.7 | 54.4 ± 18.26 | 58.4 ± 14.9 | 58.7 ± 15.8 | 58.2 ± 14.9 |
| Male | 24 (63.1%) | 3 (30%)<sup>o</sup> | 21 (75%)<sup>o</sup> | 4 (44.4%)* | 17 (89.5%)* |
| **Race %** |  |  |  |  |  |
| White | 37 (97.3%) | 10 (100 %) | 27 (96.4%) | 9 (100 %) | 18 (94.7%) |
| Mixed race: American Indian or Alaskan Native and White | 1 (2.6 %) | 0 | 1 (3.6%) | 0 | 1 (5.3%) |
| **Ethnicity** |  |  |  |  |  |
| Not Hispanic or Latino | 34 (89.4%) | 9 (90 %) | 26 (89.3%) | 7 (77.8%) | 18 (94.7%) |
| Hispanic or Latino | 3 (7.9%) | 1 (10%) | 2 (7.1%) | 1 (11.1%) | 1 (5.3%) |
| Middle Eastern | 1 (2.6%) | 0 | 1 (3.6 %) | 1 (11.1%) | 0 |

### Selectively evaluating viable skin microorganisms

Sequencing-based studies exploring the impact of CHG antiseptic exposure have had conflicting results,^21–25^ likely due to DNA from the CHG killed microbes remaining on the skin.^22,26^ To selectively evaluate DNA from live bacteria within skin microbial communities, prior to DNA extraction, samples were split, with half being treated with Propidium monoazide (PMAxx) to irreversibly bind free DNA and DNA within compromised cells (**Fig. 2A**). To evaluate the total DNA from both live and dead bacteria the remaining sample half remained untreated.

To determine the optimal PMAxx concentration for complex but low biomass skin samples, swabs of the skin microbiome were first collected from the volar forearm of healthy volunteers (**Fig. 2B**). Overall, 10 µM PMAxx performed well at accurately stratifying the higher percentage of DNA from live bacteria in the control sample (58±13%) compared to in the heat killed control (5±1%) or from skin treated with CHG antiseptic (11±3%; both p-values < 0.01 both vs. control, t-tests with Welch’s correction). The comparatively low percentage of live bacteria in the control sample treated with 50 µM PMAxx (25%) suggests that excessive PMAxx in the solution is cytotoxic.

Skin microbial bioburden and community structure often varies across skin sites. To confirm the use of 10 µM PMAxx would be sufficient for viability evaluations across skin sites, swabs were collected from the back (sebaceous site), volar forearm (dry), or umbilicus (moist). This concentration of PMAxx sufficiently differentiated the percentage of DNA from viable microbes in control conditions vs. the heat killed sample across each body site (all p-values < 0.05, t-test with Welches correction. **Fig. 2C**). Together, **Figure 2B-C** highlight the importance of identifying the appropriate PMAxx concentration for the intended study community type and contradict the recent claim that under normal circumstances the majority of bacterial DNA on the skin are from diseased microbes.^27^ Across all experiments and sampled body sites, the percentage of live bacteria on unperturbed skin ranges from 43-86% for dry body sites (95% confidence interval of 58-65%) and from 43-99% for moist sites (95% confidence interval of 70-76%; **Table S2, Fig. 2B-C, Fig. S1A).**

### Pre-surgical preparation with CHG reduces both total and viable microbial bioburden the day of surgery

To determine the impact of pre-surgical preparation with CHG on skin microbial bioburden, swabs of the skin were collected from each subject’s intended surgical site and a control site during their initial clinical visit multiple times the day of surgery, and their follow-up clinic visit (**Fig. 1B**). All subjects showered with 4% CHG soap the night before and morning of their surgery (before the Pre-OR sample collection) and 2% CHG / 70% Isopropanol was applied to just the surgical field prior to incision (before the OR sample; **Table S1**). Microbial culture is generally used to evaluate CHG efficacy.^13–18^ Swabs were collected from the surgical site during recovery in the post-operation care unit for quantitative bacterial culture. On the day of surgery, 5/28 subjects had culturable bacteria (**Fig. S1B**).

Viability-qPCR of the bacterial 16S rRNA gene was used to quantify total and viable bacterial bioburden across all timepoints (**Fig. 3A**). Due to high inter-individual variations in baseline microbial bioburden, the log_10_(fold change) from each subject’s baseline surgical site or control site bioburden was calculated (**Fig. 3A, Table S3**). Showering with CHG soap induced a roughly 1.25 log_10_ fold reduction in the viable microbiome at both moist and dry surgical sites (p-values < 0.01 pre-OR vs. baseline). This corresponded with a 0.84 log_10_ reduction in total microbial DNA (p-value < 0.01 pre-OR vs. baseline). Application of CHG to the surgical site just before incision further reduced the viable microbial bioburden an additional 0.19 log_10_ and 0.04 log at moist and dry surgical sites respectively (all p-values

**Figure 3:**
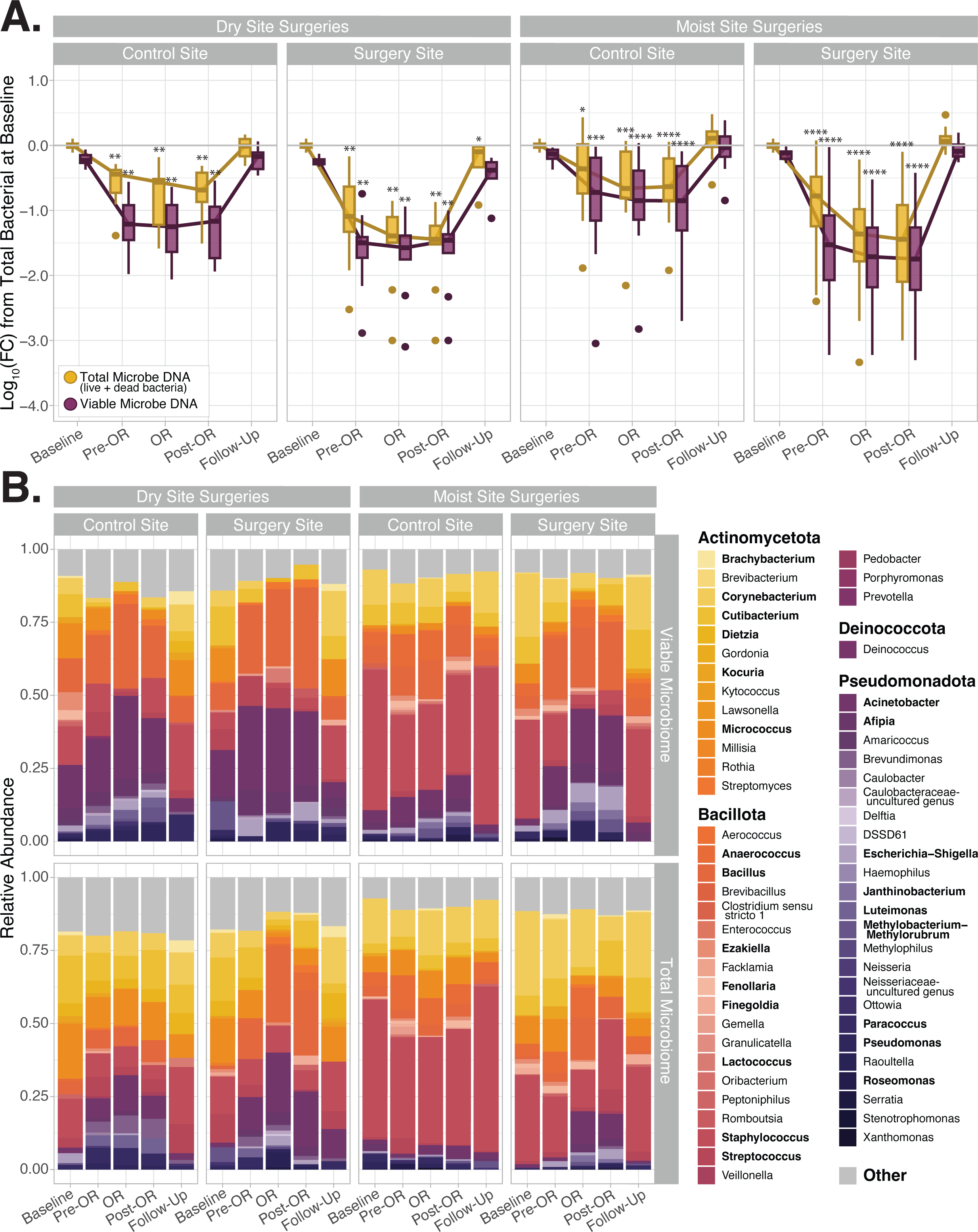
Viable and total subject skin microbial communities over time. A. Exposure to CHG induces a significant reduction in both total (yellow) and viable (purple) microbial communities the day of surgery (Pre-OR, OR, and Post-OR timepoints). Data displayed as the Log_10_(fold change) from each subject’s total bacterial bioburden at baseline. P-values represent Wilcoxon matched pairs ranked tests between the fold change in total or viable bioburden at a later timepoint vs. the respective total or viable microbial bioburden at baseline. More detail on the viable and total bioburden along with additional statistical comparisons is provided in **Supplemental Table 3**. Details on the percentage of live bacteria in these samples is in **Supplemental Figure 1. B.** Plots display the average relative abundance of each genus at the surgical or control site at each timepoint for all the subjects who underwent moist or dry surgeries. Average genera relative abundance within the viable and total (live + dead) microbial communities are shown in the top and bottom row, respectively. Bolded genera are those comprising at least 30% of the microbial community in at least one sample. Other represents taxa < 0.5% in proportion.

< 0.01 OR vs. baseline). In general, moist surgical sites displayed greater reduction in microbial communities at the OR and post-OR timepoints than dry surgical sites, likely due to greater baseline microbial bioburden. Control sites exhibited roughly 0.75 and 0.43 log_10_ fold reduction in their viable and total microbial bioburden, respectively, at the pre-OR sample collection (all p-values < 0.01), likely secondary to exposure to CHG soap during the pre-operative showers. Further details on the percentage of DNA from live bacteria at each timepoint are illustrated in **Supplemental Figure 1A.**

### Skin microbial communities before, during, and after the day of surgery

To characterize the impact of CHG antiseptic on skin microbial community structure, both viable and total bacterial DNA sample portions underwent 16S rRNA profiling. *Staphylococcus, Corynebacterium, Cutibacterium, Acinetobacter,* and *Bacillus* genera comprised the main taxa in viable and total (both live + dead bacterial DNA) communities across all sites (**Fig. 3B**). No significant differences were observed in viable or total microbial community structure in samples collected at baseline or at the post-surgical clinic follow-up (p-values > 0.05, PermANOVA, **Fig. S2A, E, Table S4**). These findings indicate that our viability assay does well in normal (baseline) conditions in selectively evaluating DNA from live microorganisms without over representing taxa from a particular phylum or genus in the viable microbiome. Viable microbial communities are, however, significantly different than the total microbiome on the day of surgery for samples collected at the Pre-OR, OR, and Post-OR timepoints (all p-values < 0.001; **Fig. S2B-D, Table S4**). These differences are largely driven by an over representation of residual DNA from *Corynebacterium*, *Micrococcus* and *Staphylococcus* within total microbial communities on the day of surgery (**Fig. S2F-H**). These findings support the hypothesis that following antiseptic application free DNA from newly killed bacteria, particularly from highly abundant skin commensal taxa, can persist on the skin.

### Subject features influencing baseline skin microbial communities

Local skin physiology and the environment can influence the skin microbiome.^7^ To accurately address the primary aim of characterizing the effects of chlorhexidine antiseptic on viable skin microbial communities, we first identified the potential confounding effects of subject and environmental factors on community composition. Bray-Curtis beta diversity was utilized to compare sample microbial community structures. Baseline viable microbial communities were then evaluated for potential associations with subject characteristics and home environment. Consistent with the well documented variation in skin microbial communities across different body sites,^6,7,33^ body site of sample collection and site characteristics such as a moist or dry environment had the largest associations with baseline skin microbial community composition (both p-values < 0.001, univariate PERMANOVA; **Fig S3A, Table S5**). Surgical and control samples taken from dry body sites (e.g., central abdomen, forearm), contained higher proportion of taxa from the *Micrococcus*, *Acinetobacter,* and *Bacillus* genera (all p-values < 0.05, FDR q-values < 0.01, = 0.03, and = 0.15 respectively; **Fig S3B-E**). Meanwhile, *Staphylococcus* dominated moist body site samples (e.g., umbilicus, armpit; p-value < 0.05, FDR q-value = 0.13, **Fig S3F**).

Baseline viable microbial communities were also significantly influenced by the subject’s gender (p-value < 0.01, univariate PERMANOVA; **Fig S3G, Table S5**). Male subjects had a significantly greater relative abundance of *Corynebacterium* and *Staphylococcus,* while female subjects had higher relative abundance of *Streptomyces,* and *Afipia* (all p-values <0.05 and, FDR q-values < 0.2; **Fig S3H-K**). Additional subject features associated with skin microbial community composition included subject BMI, an history of MRSA infection, acne, and living with a dog (p-values <= 0.05, univariate PERMANOVA; **Table S5**). However, these associations were not significant when considering gender and body site in a multivariate assessment.

### Surgical features associated with viable skin microbial community composition the day of surgery

Samples of the viable microbiome collected on the day of surgery were evaluated with Bray-Curtis beta diversity and assessed for potential associations with features related to the surgery (e.g., antibiotic prophylaxis and length of surgery, **Table S6**). Body site of sample collection and type of surgery (e.g., cholecystectomy, inguinal hernia repair) had the strongest association with viable microbial community composition the day of surgery (both p-values < 0.001, univariate PERMANOVA). Antibiotic prophylaxis with cefazolin, even after accounting for gender and moist/dry body site as confounding variables, was strongly associated with skin microbiome composition (p-value < 0.001 multivariate PERMANOVA). Patients who received antibiotic prophylaxis displayed a significantly decreased relative abundance of *Bacillus* and *Streptococcus* species at their surgical sites and a corresponding increase in *Pseudomonas* and *Afipia* compared to those that did not receive antibiotics (p-values < 0.05, FDR corrected q-values = 0.06, 0.13, 0.19, and 0.19 respectively, **Fig. S4A-D**).

### Pre-surgical preparation with CHG promotes temporary shifts in the skin microbiome

The longitudinal study design allowed us track how the microbiome recovers following CHG application. After accounting for body site, gender, and antibiotic prophylaxis, the viable skin microbial community compositions at the surgical site on the day of surgery were significantly different than at the baseline or follow-up clinical visit (p-value < 0.0001, multivariate PERMANOVA; **Table S7**). With univariate assessment, the timepoint of sample collection had the strongest association with viable community structure at the surgical site (p-value < 0.0001, univariate PERMANOVA; **Fig. 4A, Table S7**), followed by the body site of sample collection (p-value < 0.001; **Fig. S5A**).

**Figure 4:**
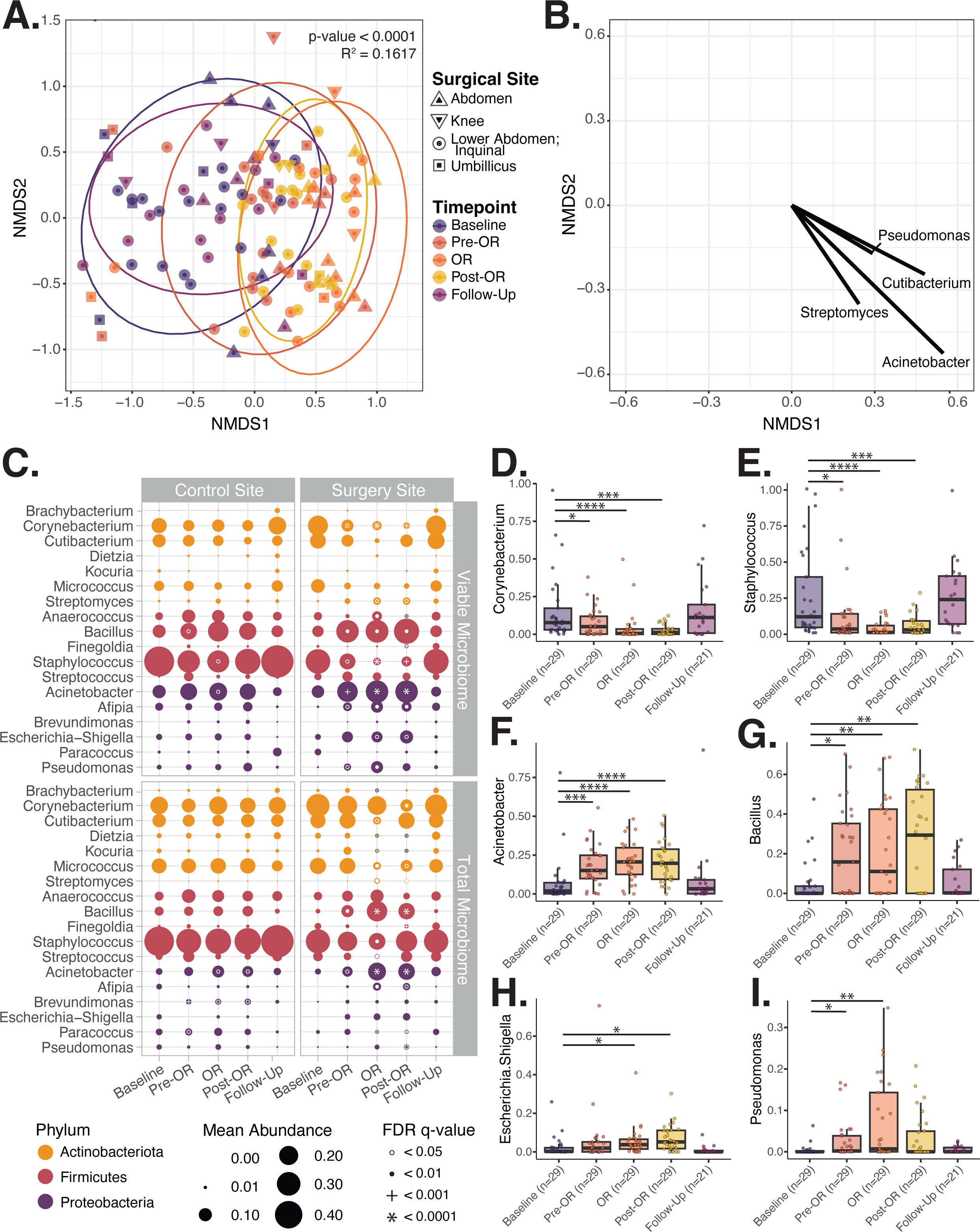
Pre-surgical application of chlorhexidine gluconate (CHG) antiseptic is associated changes in viable microbial community composition, particularly at the surgical site, on the day of surgery. A. Bray-Curtis beta-diversity Non-metric MultiDimensional Scaling (NMDS) ordination displaying the significant shift in viable microbial community composition at subject surgery sites over time. Difference in microbial community composition at each time point assessed via PermANOVA with 9999 permutations. **B.** Vector plot displaying the taxa whose relative abundance significantly correlates with a points position on the NMDS plot. Only taxa with a spearman correlation p-value < 0.05 are shown. The longer the vector the larger the spearman rho. **C.** Plot displaying the change in the average relative abundance of key taxa in the viable and total microbiome at control and surgical sites over time. Average taxa abundance across samples is indicated by the size of the point. Differential relative abundance of taxa at later timepoints compared to baseline was evaluated via MAASLIN2 accounting for the individual subject, subject gender, body site of sample collection, and use of pre-surgical antibiotic prophylaxis as random effects. White or grey circles, solid dots, plus sign, and asterixis indicate the degree of significance. **D-I.** Relative abundance plots illustrating significant changes in key taxa within viable microbial communities at both moist and dry surgical sites over time. Points indicate the relative abundance of that taxa within a specific subject’s sample. Note; one subject underwent simultaneous umbilical and inguinal hernia repair. Both sites were sampled at all timepoints. Thus, the n = 29 at the baseline through Post-OR timepoints and n=21 at follow-up, which is one more than the total number of subjects who underwent surgery and came for in-person follow-up, n = 28 and = 20 respectively.

On the day of surgery, surgical sites were enriched in the proportion of several bacterial genera associated with SSI, including*, Acinetobacter, Bacillus, Escherichia-Shigella,* and *Pseudomonas*, compared to baseline communities (all FDR q-values < 0.05, Pre-OR, OR or Post-OR vs, Baseline, **Fig. 4B-I, Fig. S5B-F, Table S8**). These changes are accompanied by a corresponding reduction in commensal taxa relative abundance, particularly *Corynebacterium, Micrococcus,* and *Staphylococcus* (all FDR q-values < 0.05, Pre-OR,OR or Post-OR vs, Baseline). These trends were consistent across dry and moist body sites (**Fig. S5, Fig S6, Table S8**). Since *S. aureus* and *S. epidermidis* are notable SSI associated pathogens, *Staphylococcus* ASV’s were assigned probable species level designation via alignment to the BLAST database. This revealed that the reduction of *Staphylococcus* species on the day of surgery corresponded to reduced coagulase negative staphylococci, particularly *S. epidermidis* (**Fig. S5G**). The increased relative abundance of *Acinetobacter* and *Bacillus* taxa on the day of surgery were primarily due to greater *A. lwoffii* and *B. subtillis* respectively (**Fig. S5H**). While *E. coli* and unclassified *Escherichia-shigella* spp. as well as *P. aeruginosa* and unclassified *Pseudomonas* spp. were the main contributors to increased *Escherichia-shigella* and *Pseudomonas* abundance.

Both viable and total surgical site microbial communities from the baseline and follow-up clinic visits did not differ (p-value > 0.05, univariate PERMANOVA, **Table S7**). Collectively, the findings displayed in **Figure 4** and **Supplemental Figure 6** demonstrate that pre-surgical preparation with CHG promotes significant shifts in microbial community composition at the surgical site. Both control and surgical site communities return to near baseline after the patient returns home (p-value > 0.05, multivariate PERMANOVA, baseline vs. follow-up**; Fig4A, Table S7, Fig S7A**). However, at follow-up, subjects’ skin microbial communities are more similar to other individual baseline microbiomes than their own (p-values < 0.01, Wilcoxon paired signed rank test, **Fig S7B**).

Total microbial communities (DNA from both live and dead bacteria), at the surgical site did demonstrate changes in community composition on the day of surgery (p-value < 0.0001, multivariate PERMANOVA; **Table S7**). However, compared to the viable microbiome, the shifts in total microbiome community structure and taxa relative abundance were less pronounced and/or delayed and not noted until the OR or Post-OR sample collection (**Fig. 4C, Fig. S6, Tables S7-8).** Changes in the relative abundance of critical taxa noted in the viable community, including increases in *Escherichia* and *Pseudomonas*, were not detected when evaluating the total microbiome (**Fig. 4C**). These findings suggest if viability had not been assessed, overrepresentation of free DNA from abundant, lysed skin commensal taxa can mask true community shifts occuring.

At control sites, both viable and total skin microbial communities were more similar over time (both p-values > 0.05, univariate PERMANOVAs; **Fig. S8A, FigS6, Table S7**) and body site of sample collection (forearm or armpit) maintained the strongest association with microbial community composition (p-values < 0.0001, univariate PERMANOVAs; **Fig. S8B, Table S7**). Nevertheless, on the day of surgery control sites had subtle changes in taxonomic composition, including a minor rise of *Acinetobacter* and *Bacillus* relative abundance and minor decline in *Staphylococcus* (all FDR q-values < 0.05, OR, Pre-OR, and OR vs. Baseline respectively; **Fig S8C-E**). Additionally, there were no significant changes in the Shannon alpha diversity of total or viable bacteria within control or surgical site samples over the time course of sample collection (**Fig. S9**).

### *Bacillus* spp. and *Enterococcus* spp. isolated from surgical patients display increased resistance to CHG

The minimum inhibitory concentration of CHG was tested against representative isolates obtained from baseline samples including representatives from the *Bacillus, Corynebacterium, Enterococcus, Micrococcus, Staphylococcus,* and *Pseudomonas* genera. Consistent with the shifts in microbial community composition, < 200 µg/ml of CHG was sufficient to prevent the growth of *Corynebacterium, Cutibacterium, Micrococcus* and *Staphylococcus* (including *S. aureus* and *CoNS Staphylococci*) species (**Table 2).** However, isolates of *Bacillus* and *Enterococcus* displayed increased resistance to CHG, requiring 250-1250 µg/ml CHG to inhibit their growth. In short, most individuals likely have some degree of antiseptic resistance within their skin microbial communities, and some may harbor highly resistant taxa even prior to antiseptic exposure (**Table 2**).

**Table 2:** *Bacillus* and *Enterococcus* isolates display increased resistance to CHG. Bacterial Isolates collected from surgical subjects were identified via Sanger sequencing of the 16S rRNA gene. The minimum concentration of CHG needed to inhibit isolate growth was measured via a disc diffusion MIC assay. Isolates were tested in duplicate and the average MIC was calculated. This table also includes the subject and sample from which each isolate was collected.

| Isolate | LK-ID | Subject | Collection Timepoint | Site | Average MIC (µg/ml) |
| --- | --- | --- | --- | --- | --- |
| *Bacillus safensis* | LK2236 | CHG-002 | Baseline | Control Site | 250 |
| *Bacillus* spp. 1 | LK3019 | CHG-025 | Baseline | Control Site | 500 |
| *Bacillus* spp. 2 | LK3063 | CHG-028 | Baseline | Control Site | 1250 |
| *Corynebacterium mycetoides* | LK2860 | CHG-021 | Baseline | Control Site | 31 |
| *Corynebacterium sanguinis* 1 | LK2319 | CHG-007 | Baseline | Control Site | 12 |
| *Corynebacterium sanguinis* 2 | LK2685 | CHG-017 | Baseline | Control Site | 47 |
| *Corynebacterium striatum* | LK2376 | CHG-003 | Follow-Up | Surgical Site | 63 |
| *Corynebacterium ureicelerivorans* | LK3068 | CHG-028 | Baseline | Control Site | 31 |
| *Cutibacterium* spp. | LK2346 | CHG-010 | Baseline | Control Site | 47 |
| *Dietzia* sp. | LK2320 | CHG-008 | Baseline | Control Site | 16 |
| *Enterococcus faecalis* | LK2699 | CHG-006 | Follow-Up | Surgical Site | 750 |
| *Micrococcus endophyticus* | LK2945 | CHG-018 | Baseline | Control Site | 16 |
| *Micrococcus luteus* 1 | LK2435 | CHG-012 | Baseline | Surgical Site | 63 |
| *Micrococcus luteus* 2 | LK2473 | CHG-014 | Baseline | Control Site | 31 |
| *Micrococcus luteus* 3 | LK2281 | CHG-005 | Baseline | Control Site | 125 |
| *Micrococcus yunnanensis* 1 | LK2439 | CHG-012 | Baseline | Surgical Site | 63 |
| *Micrococcus yunnanensis* 2 | LK2686 | CHG-017 | Baseline | Control Site | 24 |
| *Pseudomonas luteola* | LK2294 | CHG-007 | Baseline | Control Site | 125 |
| *Staphylococcus aureus* | LK2886 | CHG-016 | Follow-Up | Control Site | 94 |
| *Staphylococcus capitis* | LK2681 | CHG-016 | Baseline | Control Site | 63 |
| *Staphylococcus epidermidis* 1 | LK2219 | CHG-001 | Baseline | Control Site | 187.5 |
| *Staphylococcus epidermidis* 2 | LK2279 | CHG-006 | Baseline | Control Site | 73 |
| *Staphylococcus hominis* 1 | LK2452 | CHG-012 | Baseline | Surgical Site | 94 |
| *Staphylococcus hominis* 2 | LK2226 | CHG-003 | Baseline | Control Site | 94 |
| *Staphylococcus* spp. 1 | LK2887 | CHG-016 | Follow-Up | Control Site | 146 |
| *Staphylococcus warneri* 1 | LK2474 | CHG-012 | Baseline | Surgical Site | 47 |
| *Staphylococcus warneri* 2 | LK2894 | CHG-016 | Baseline | Control Site | 47 |

## Discussion

Surgical site infections continue to occur despite the widespread adoption of surgical antiseptics, including CHG. Despite limited evidence, the potential for large scale impacts on the healthy skin microbiome is enormous and remains understudied. This study characterizes the immediate and long-term effects of pre-surgical preparation with CHG on the absolute burden and community structure of viable skin-associated bacteria in patients undergoing elective surgeries (**Fig. 1**). Our findings demonstrate that pre-operative preparation with CHG i) effectively reduces viable microbial bioburden at the surgical site and ii) promotes a microbial community composition enriched for several potentially pathogenic taxa, particularly Gram-negative bacterial taxa. We also demonstrate that most individuals’ microbiomes recover within 2-4 weeks post-perturbation. This indicates that CHG induced changes are transient and the skin microbiome itself is resilient to these insults. However, we also observed that patients can harbor CHG resistant taxa on their skin, even before surgery. In short, these findings underscore that broad scale application of topical antiseptics like CHG may disrupt healthy skin microbiota resulting in space for others to proliferate, particularly pathogenic Gram-negative and biofilm forming taxa.

Chlorhexidine gluconate has a unique ability to bind bacterial DNA to the skin surface.^22^ This property is likely why sequencing based studies on the effects of CHG on skin microbial communities have had inconsistent findings.^21–25^ To circumvent DNA from newly killed bacteria confounding sequencing results, we optimized a propidium monoazide (PMAxx) based viability assay for selective evaluation of DNA from live microorganisms within the skin microbiome (**Fig. 2**). After determining the ideal sample collection solution and PMAxx concentration (10 µM), we show that under normal circumstances approximately 60-80% of bacterial DNA on the skin is from viable microbes within the community (**Fig. 2B-C, Table S2**). This stands in contrast to the recent findings reported by Acosta et al. that bacterial DNA on the skin surface overrepresents the viable skin microbiome.^27^ Once activated PMA and its close derivative PMAxx are highly reactive nitrene compounds that irreversibly bind to double stranded DNA and prevent it from being amplified or sequenced.^34,35^ However, too high a concentration of PMA or PMAxx for the sample biomass can result in false negatives, either due to being toxic to live cells or PMA/PMAxx remaining active beyond the duration of photoactivation and subsequent interaction with DNA released during the extraction process from live cells.^35^ This, in tandem with our observation that excessive (50 µM) PMAxx negatively interacts with the low biomass of skin microbial communities (**Fig. 2B**), we believe that Acosta et al.’s use of 50 µM PMA was cytotoxic, resulting in a misrepresentation of the proportion of DNA from dead microbes on the skin.

There is always a concern that incorporation of a new method may bias sequencing results to overrepresent a particular class of microbes.^36^ With this in mind we compared subjects’ viable (live microbial DNA) to their total (both live and dead microbial DNA) microbial community compositions (**Fig. S2A, E**). The similarity of patients’ viable and total microbiomes at baseline and their clinical follow-up confirm that our viability protocol works well in normal, baseline conditions selectively evaluating DNA from live microbes without overrepresenting groups of taxa from skin microbial communities.

Significant differences between the viable and total microbial bioburden and community composition on the day of surgery (**Fig. S2B-D, Table S3**) are consistent with the observation that CHG likely binds to and retains microbial DNA at the skin surface.^22^ On the day of surgery, total skin microbial communities also contained a higher relative abundance of several commensal skin taxa, *Corynebacterium, Micrococcus* and *Staphylococcus,* compared to viable communities (**Fig. S2F-H**). Changes in the relative abundance of critical taxa noted in the viable community at the surgical site after exposure to CHG were also missed when evaluating the total microbiome (**Fig. 4C**). Together these findings illustrate how following CHG application free DNA from newly killed bacteria, particularly from highly abundant skin commensal taxa, likely persists on the skin and can subsequently prevent sequencing-based studies from identifying the true microbial community shifts induced by antiseptic exposure. This further underscores the importance of assessing the viable portion of the microbiome in this and similar contexts of antiseptic, antibiotic, or other microbially toxic exposure studies.

Importantly, with our optimized viability assay, we report that CHG antiseptic does indeed impact the skin microbiome (**Fig. 3**, **Fig. 4**). We observed that pre-surgical application of CHG significantly reduces viable microbial bioburden (**Fig. 3A**). However, complete sterility is not achieved, with roughly 80-200 live bacteria per inch^2^ skin remaining at the surgical site on the day of surgery (**Fig. 3A, Table S3**). This both reinforces the efficacy of CHG shown in culture-based studies, but, also contradicts the sterility endorsed by these culture-based validations.^13–18^

Pre-surgical preparation with CHG also influenced viable microbial community compositions at the surgical site (**Fig. 4, Table S7**). After accounting for key factors known to influence skin microbial community composition (e.g., gender, body site of sample collection, surgical antibiotic prophylaxis), viable microbial communities display reduced relative abundance of several commensal skin taxa, particularly *Corynebacterium, Micrococcus,* and coagulase negative *Staphylococcus* (CoNS) species at the surgical site on the day of surgery (**Fig. 4C-E, Fig. S5C-E, Table S8**). Although generally regarded as beneficial,^37,38^ *S. epidermidis* can be an opportunistic pathogen and is one of the leading causes of SSI, particularly for surgeries involving implanted devices (eg. mesh or joint replacements).^1,39^ Lower relative abundance of CoNS, specifically *S. epidermidis,* on the day of surgery indicates that CHG antisepsis and current pre-surgical protocols target this opportunistic pathogen well (**Fig. S5G**). SSI infections are typically caused when bacteria from the patient’s endogenous flora are inoculated into the surgical site at the time of surgery, as highlighted by SSI due to *S. epidermidis*.^1,40^ Skin commensals help maintain community homeostasis in part through microbial-microbial interactions that prevent pathogen overgrowth and prompt pathogens to maintain less-virulent dispositions.^7,41–45^ The relative loss of numerous commensal taxa following CHG (**Fig 4. Fig S5**) could also result in a skin microbiome that is unable to properly defend against opportunistic pathogens.

Baseline microbial community compositions with subtle patterns of antiseptic resistance likely influence community dynamics following pre-surgical preparations. Some individuals harbor highly resistant taxa prior to pre-surgical antiseptic exposure (**Table 2**). Overall our findings are consistent with multiple reports that CHG is less effective against Gram-negative bacteria and microbes within a biofilm.^14,21,49–51^ *Staphylococcus aureus, S. epidermidis, Escherichia coli,* and *Pseudomonas aeruginosa* are among the most common causes of SSI.^1^ Although not significant, *S. aureus* ASVs were also in higher relative abundance at the OR sample collection (**Fig. S5G**). The incomplete efficacy of CHG against these taxa may partially explain why these pathogens continue to commonly cause SSI.

Reassuringly, most subjects’ viable microbial bioburden and community composition resembled baseline at their follow-up clinic visit (**Fig. 3A**, **Fig. 4, Fig. S5H, Fig. S7, Table S8**). However, these skin microbial communities were more similar to other individuals baseline microbiome composition than their own (**Fig S7B**). This suggests that application of CHG may in a way reset the microbiome. Although we were unable to interrogate precisely when skin microbial communities repopulate, our data suggest they start to recover within hours. For instance, although their duration of activity is shorter,^54^ bacterial bioburden and microbiome diversity can recover as early 6 hours after ethanol or povidone-iodine application.^22,55^ Sources of repopulation likely include microbes residing deep within hair follicles,^56^ neighboring body sites that received a lower dose of CHG, as well as patients’ own clothes and home environment.^7,57^ Together, these findings reinforce the general resiliency of skin microbial communities to acute perturbations and support that for individuals preparing for surgery, the effects of CHG antiseptic on skin microbial communities are temporary.

One of the main limitations of this study was that we only enrolled subjects undergoing elective outpatient surgeries. All subjects went home the same day of their surgery preventing an assessment of microbiome changes in the first few days post-application. To address this, future works aim to expand this study design to more surgery types, particularly surgeries requiring post-operative inpatient stays. Another limitation is the taxonomic resolution afforded by 16S rRNA gene sequencing. Metagenomic shotgun sequencing would provide a more comprehensive insight into strain-level, multi-kingdom, and functional impacts of CHG on the skin.

In conclusion, we confirm that pre-surgical application of CHG does reduce viable skin microbial bioburden. However, complete sterility is not achieved. Chlorhexidine induces a temporary shift in microbial composition. Skin microbial communities at the time of surgery are enriched for several potentially pathogenic taxa, notably *Pseudomonas, Escherichia, Acinetobacter,* and *Bacillus*. Incomplete antiseptic efficacy against these taxa likely places some patients at disproportionate risk for developing an SSI. Importantly, with this work we’ve addressed a critical uncertainty in the field.^21–25^ To our knowledge, this is the first work to accurately demonstrate the impact of CHG on viable, rather than total, skin microbial bioburden and community composition via sequencing-based methods. Collectively, our findings identify tangible avenues to improve antiseptic formulations and pre-surgical preparation protocols to better target Gram-negative and biofilm forming microbes while protecting beneficial skin commensals. With these efforts we can further reduce the burden of surgical site infections.

## Acknowledgements

We also would like to acknowledge and extend special thanks to Derek Gonzalez, Sophie Oubaha, and the University of Wisconsin Department of Surgery Clinical Research Team for their assistance in with subject recruitment, enrollment, sample collection, and managing this study.

Additional thanks to Alison Jayme, NP, the UW-Health General Surgery Clinic nurses and staff, and all members of the operating room teams for their assistance with this study and patient care.

Special thanks to J.Z. Alex Cheong, PhD, for his guidance on the methodology and assistance with sample collection.

We also thank the University of Wisconsin Biotechnology Center for their expertise and microbial sequencing.

## Contributions

**ECT:** conceptualization, methodology, formal analysis, investigation, data curation, visualization, writing – original draft, writing-review & editing. **KX:** methodology, validation, formal analysis, investigation, data curation, visualization, writing-review & editing. **KDLC:** methodology, investigation, data curation, writing-review & editing. **LH:** methodology, investigation, data curation, writing-review & editing. **SS:** methodology, investigation, data curation, writing-review & editing. **DA:** investigation writing-review & editing. **OG:** investigation, writing-review & editing. **JS:** supervision, resources, writing-review & editing. **AH:** supervision, resources, writing-review & editing. **AG:** conceptualization, supervision, resources, funding acquisition, writing – original draft, writing-review & editing. **LRK:** conceptualization, supervision, resources, funding acquisition, writing – original draft, writing-review & editing.

## Supplemental Figure Legends

**Supplemental Figure 1: Viable Microbial bioburden. A.** Companion figure to Figure 3A. Points represent the percent live bacteria within each subject’s control and surgical site samples over time. Data also grouped by whether the sample was from a moist or dry body site. **B.** Swabs of the skin microbiome at subjects’ surgical sites in the post-operative care unit and from both the control and surgical site during the post-surgery clinic visit. Plot displays the culturable bacterial bioburden at moist and dry sampling sites at both timepoints.

**Supplemental Figure 2: Viable and total microbial community compositions differ on the day of surgery. A-E.** Non-metric Multidimensional Scaling (NMDS) ordination of the Bray-Curtis beta-diversity at each timepoint. PERMANOVAs with 9999 permutations were utilized to evaluate the differences between the viable (PMAxx treated) and total (not treated) sample community compositions. Details can be found in **supplemental table 4. F-H.** MAASLIN2 was used to determine differences in the relative abundance of individual taxa between viable and total communities from samples collected on the day of surgery (Pre-OR, OR, and Post-OR timepoints combined). **** indicates FDR q-value with Benjamini-Hochberg correction < 0.0001.

**Supplemental Figure 3: Microbial community composition at baseline is associated with body site of sample collection and subject gender. A.** NMDS ordination of Bray-Curtis beta-diversity of viable microbial communities at subjects’ baseline clinic visit highlighting the association between the microbiome composition and body site of sample collection (univariate PERMANOVA with 9999 permutations). **B.** plot of average relative abundance of key genera within viable (top row) and total (bottom row) microbial communities across body sites samples collected at the baseline timepoint. Size of the dot indicates the average genera relative abundance. **C-F.** MAASLIN2 was used to determine differences in the relative abundance of individual taxa within viable communities of samples collected at moist body sites (armpit, lower abdomen to inguinal, and umbilicus combined) compared to those from dry body sites (forearm, central abdomen, and knee combined). Both control and surgical site samples were included in this analysis. In these calculations both subject and gender were incorporated as random effects. Only taxa with significantly different relative abundance (p-value < 0.05) between the groups are shown. FDR q-values with Benjamini-Hochberg correction indicated. **G.** Bray-Curtis beta diversity NMDS ordination highlighting the association of subject gender with baseline viable microbial community composition (univariate PERMANOVA with 9999 permutations). **H-K.** Differences in the relative abundance of individual taxa within the viable baseline microbiome of male and female subjects were assessed via MAASLIN2. Both control and surgical site samples were included in this analysis and subject and body site of sample collection were incorporated as random effects into these calculations. Only taxa with significantly different relative abundance (p-value < 0.05) between the groups are shown. FDR q-values with Benjamini-Hochberg correction indicated.

• FDR q-value < 0.2

* FDR q-value < 0.05

** FDR q-value < 0.01

*** FDR q-value < 0.001

**** FDR q-value < 0.0001

**Supplemental Figure 4: Antibiotic prophylaxis with cefazolin is associated with reduced relative abundance of *Bacillus* and *Streptococcus* and increased *Afipia* and *Pseudomonas* at the surgical site on the day of surgery.** A-D. Viable microbiome samples from the surgical site on the day of surgery, the samples collected at the pre-OR, OR, and post-OR timepoints, were assessed collectively. Differences in taxa relative abundance were evaluated with MAASLIN2 incorporating the individual subject, gender, and body site of sample collection as random-effects. Only taxa with significantly different relative abundance (p-value < 0.05) between the groups are shown. FDR q-values with Benjamini-Hochberg correction indicated.

**Supplemental Figure 5: Exposure to Chlorhexidine Gluconate is associated with changes in viable microbial community composition at the surgical site on the day of surgery.** Continuation of Figure 4**. A.** Bray-Curtis beta-diversity NMDS ordination highlighting the association between viable microbial community composition and the body location for the surgical site (univariate PERMANOVA with 9999 permutations). Details can be found in **supplemental table 7. B-F.** Relative abundance plots for taxa significantly more or less abundant in the viable microbiome at the surgery site on the day of surgery, after exposure to CHG, compared to the baseline sample collection. Differential abundance of taxa at each later timepoint compared to the baseline timepoint were assessed via MAASLIN2 and evaluations were made accounting for the individual subject, subject gender, body site of sample collection, and antibiotic prophylaxis as random effects. **Supplemental table 8** contains further details. Note; one subject underwent simultaneous umbilical and inguinal hernia repair. Both sites were sampled at all timepoints. Thus the n = 29 at the baseline through Post-OR timepoints and n=21 at follow-up, which is one more than the total number of subjects who underwent surgery and came for in-person follow-up, n = 28 and = 20 respectively. **G.** ASVs from the *Staphylococcus* genus were aligned against the BLAST database to obtain probable species assignment. Plot of mean relative abundance of *Staphylococcus* species within viable microbial communities over time. Apart from *S. aureus,* all species identified are coagulase negative *Staphylococci* (CoNS). “CoNS *Staphylococcus* spp.” indicates ASVs that aligned well to several CoNS species. **H.** ASVs from the *Acinetobacter, Bacillus, Escherichia-shigella,* and *Pseudomonas* genera were aligned against the BLAST database to obtain probable species assignment. Plot displays mean relative abundance of these species within viable microbial communities over time.

**Supplemental Figure 6: Change in average relative abundance of key taxa over time at surgical and control sites collected from most and dry body sites.** Companion to Figure 4C displaying the average relative abundance of each taxa over time in the viable and total microbial communities at moist and dry surgical sites and control sites respectively. Mean taxa abundance is indicated by the size of the point. Differential relative abundance of taxa at later timepoints versus at baseline was evaluated via MAASLIN2. All MAASLIN evaluations were made accounting for subject, body site of sample collection, gender, and antibiotic prophylaxis as random effects. White, or in a few cases grey, circles, filled-in dots, plus sign, and asterixis indicate the degree of significance. Further details for the MAASLIN2 results located in **Supplemental Table 8.**

**Supplemental Figure 7: Change microbial community composition from baseline. A.** Similarity of each subject’s surgical or control site microbiome at each timepoint was compared to their respective baseline community composition via the Bray-Curtis beta diversity metric. **B.** Bray Curtis beta diversity metric was also used to evaluate the similarity of each subject’s microbiome composition at follow-up to both their own baseline microbiome composition and the most similar baseline microbial community composition (smallest Bray-Curtis distance) of another subject. Differences between the bray-curtis distance between a subject’s follow-up to their own baseline versus someone elses baseline microbiome were evaluated with the Wilcoxon matched pairs signed rank tests. ** indicates p-value < 0.01; **** indicates p-value < 0.0001.

**Supplemental Figure 8: Viable communities at the control sites are more associated with the body site of sample collection**. Companion figure to figure 4 and **supplemental figure 5. A.** Bray-Curtis beta diversity NMDS ordination illustrating the strong association of body site of control samples with viable microbial community composition. **B.** Bray-Curtis beta-diversity ordination highlighting the lack of association between viable microbial community composition and timepoint of control site sample collection. Associations of microbial community composition with various features were evaluated via univariate PERMANOVAs with 9999 permutations. Details can be found in **supplemental table 7. C-E.** Relative abundance of *Acinetobacter* (**C**)*, Bacillus* (**D**), and *Staphylococcus* (**E**) in viable communities at both moist (armpit) and dry (forearm) control sites over time. Differential abundance of taxa at each timepoint compared to the baseline timepoint were assessed via MAASLIN2. All MAASLIN2 evaluations were made accounting for individual subjects, gender, body site of sample collection and antibiotic prophylaxis as random effects.

**Supplemental Figure 9: No change in Shannon alpha diversity following CHG application.** Shannon alpha diversity metric was used to measure the microbial diversity within each sample. Boxplots represent the Shannon index distribution (median ± interquartile range) of viable and total microbial communities at the surgical and control sites. Samples are grouped by if they were collected at a moist or dry body site.

## Supplemental Table Legends

**Supplemental Table 1: Summary of subjects’ home and work environment, co-morbidities, and details related to the surgery and pre-surgical prep.** Information related to pre-surgical assessments, surgery, post-surgical care, following post-operative outpatient evaluations and patient co-morbidities were extracted from the medical record. Here, significant alcohol use is defined as a recorded history of alcoholism or consuming 14 or more drinks per week at the time of enrollment. All patients who received antibiotic prophylaxis received cefazolin. Subjects also completed a survey to collect information on their skin health, hygiene habits, and home and work environments. Significant differences in between the subjects who underwent surgeries at dry versus moist body sites were determined by either Mann-Whitney or Fishers exact test.

∘ p-value < 0.05 Mann-Whitney test

* p-value < 0.05 Fishers exact test

** p-value < 0.01 Fishers exact test

*** p-value < 0.001 Fishers exact test

**Supplemental Table 2: Percentage of live bacteria at unperturbed dry, moist, and sebaceous skin sites across experiments.** Total (live and dead) and viable bacterial bioburden was determined via viability-qPCR. Percent live bacteria represents the calculated viable bacteria divided by the calculated total bacteria. For consistency, only samples processed with 10 µM PMAxx from the optimization experiment are included. For this surgical study, only control site and surgical site samples collected at the subjects’ baseline and post-surgery clinic visit are included. Data represented as mean ± standard deviation (n = the number of samples included from each experiment).

**Supplemental Table 3: Viable and total microbial bioburden at both control and surgical sites over time.** Viable and total bacterial bioburden were determined via viability-qPCR. First two columns display the median (interquartile range) of the log_10_(viable or total bacteria). Total and viable bacterial burden at each timepoint were compared via Wilcoxon matched pairs signed rank tests (small circles indicate p-values). The remaining columns show the median (interquartile range) of each subjects’ log_10_(fold change) of viable bacteria at a given timepoint from their viable bacteria or total bacteria at the baseline timepoint, third and fourth column respectively, or the log_10_(fold change) of total bacteria from the total bacteria at baseline, final column. Comparisons between bacterial bioburden or the fold change at one timepoint to the respective baseline were made also with the Wilcoxon matched pairs signed rank tests (asterisk indicate p-values).

∘ p-value < 0.05 Total v. Viable at same timepoint

∘∘ p-value < 0.01 Total v. Viable at same timepoint

∘∘∘ p-value < 0.001 Total v. Viable at same timepoint

∘∘∘∘ p-value < 0.0001 Total v. Viable at same timepoint

* p-value < 0.05 v. respective baseline microbiome

** p-value < 0.01 v. respective baseline microbiome

*** p-value < 0.001 v. respective baseline microbiome

**** p-value < 0.0001 v. respective baseline microbiome

**Supplemental Table 4. Viable and total microbial community compositions differ on the day of surgery.** Univariate PERMANOVAs with 9999 permutations were utilized to evaluate the differences between the viable (PMAxx treated, containing only DNA from viable bacteria) and total (not treated, contaiinig DNA from both viable and dead bacteria) microbial community compositions at each timepoint and all timepoints combined. Corresponding NMDS plots are in **supplemental figure 2**.

**Supplemental Table 5: Body site of sample collection and subject gender have the strongest association with the baseline viable microbiome.** Similarity of baseline viable microbial communities were assessed with Bray-Curtis beta diversity. Association between subject factors with microbiome composition were assessed both with univariate PERMANOVAs and multivariate PERMANOVAS accounting for subject gender and body site of sample collection.

**Supplemental Table 6: On the day of surgery, viable microbial community composition is associated with body site of sample collection, the type of surgery, and if the subject received antibiotic prophylaxis.** Similarity of baseline viable microbial communities were assessed with Bray-Curtis beta diversity. Association between features of the surgery and sample collection with microbial community composition were assessed both with univariate PERMANOVAs and multivariate PERMANOVAS accounting for subject gender and body site of sample collection.

**Supplemental Table 7: Pre-surgical use of chlorhexidine gluconate (CHG) antiseptic is associated with shifts in microbial community composition, particularly at the surgical site.** Swabs of the skin microbiome were collected at subject’s initial clinical visit (Baseline), multiple times the day of surgery, and their follow-up post-surgical clinic visit. All subjects showered with 4% CHG soap both the night before and morning of their surgery (before the Pre-OR sample collection) and 2% CHG / 70% Isopropanol was applied to just the surgical field prior to incision (before the OR sample). Similarity of microbial communities at the surgical and control sites were assessed with Bray-Curtis beta diversity. Associations between timepoint of sample collection, as a proxy for CHG exposure, with the composition of microbial communities were assessed both with univariate PERMANOVAs and multivariate PERMANOVAS accounting for subject gender, body site of sample collection and whether the subject received antibiotic prophylaxis.

**Supplemental table 8: Significant changes in taxa relative abundance at surgical and control sites compared to baseline.** This table is a companion to Figure 4 and **Supplemental Figures 5 & 6.** MAASLIN2 was used to evaluate differential relative abundance of taxa within viable and total surgical site and control site microbiomes at later timepoints compared to baseline. All analyses were conducted incorporating the individual subject, subject gender, body site of sample collection, and use of pre-surgical antibiotic prophylaxis as random effects.

