## Supplemental Figures for "Still Not Sterile: Chlorhexidine gluconate treatment does not completely reduce skin microbial bioburden and promotes pathogen overabundance in patients undergoing elective surgeries"

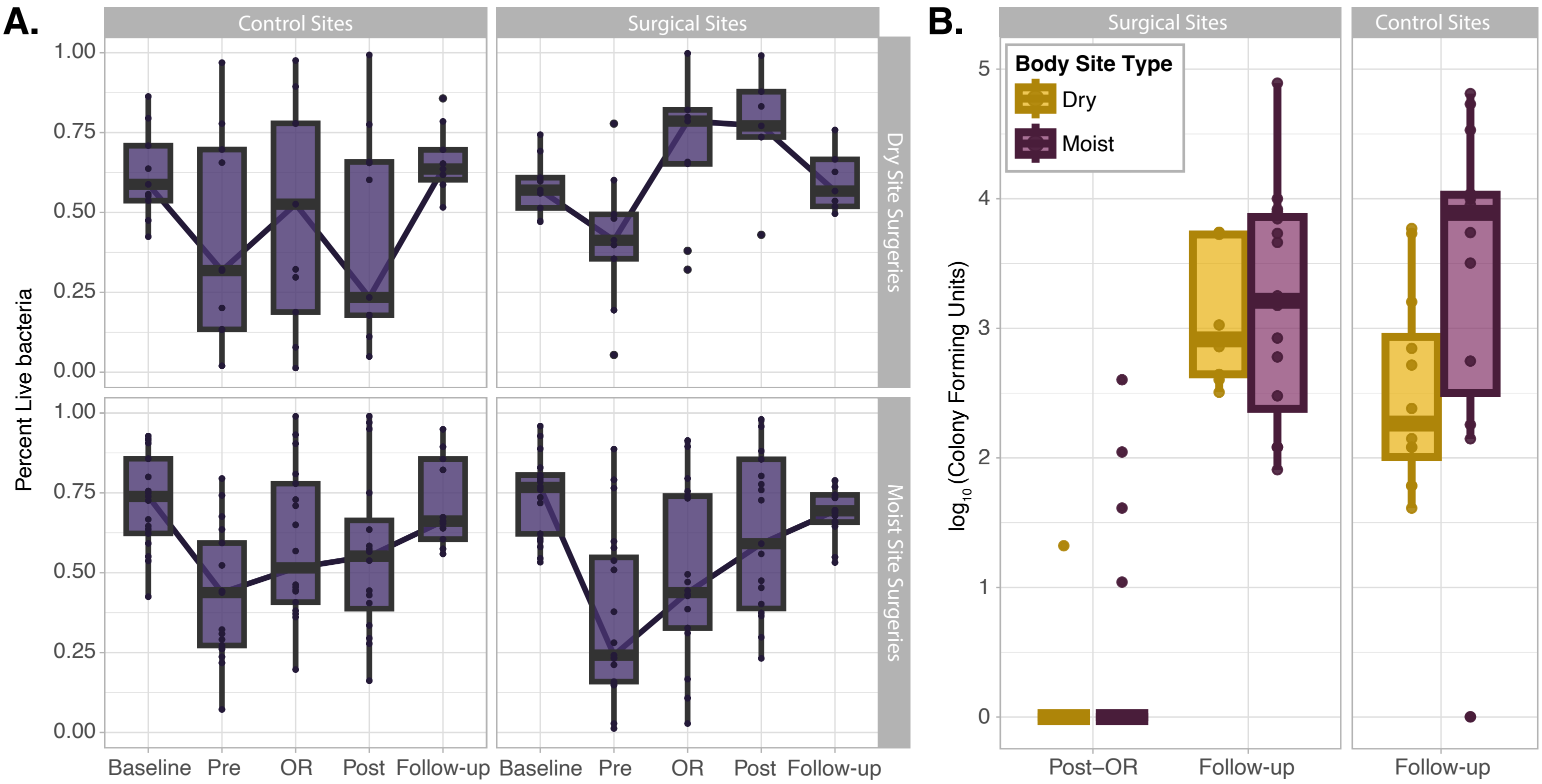


**Supplemental Figure 1: Viable Microbial bioburden. A.** Companion figure to **Figure 3A.** Points represent the percent live bacteria within each subject's control and surgical site samples over time. Data also grouped by whether the sample was from a moist or dry body site. **B**. Swabs of the skin microbiome at subjects' surgical sites in the post-operative care unit and from both the control and surgical site during the post-surgery clinic visit. Plot displays the culturable bacterial bioburden at moist and dry sampling sites at both timepoints.

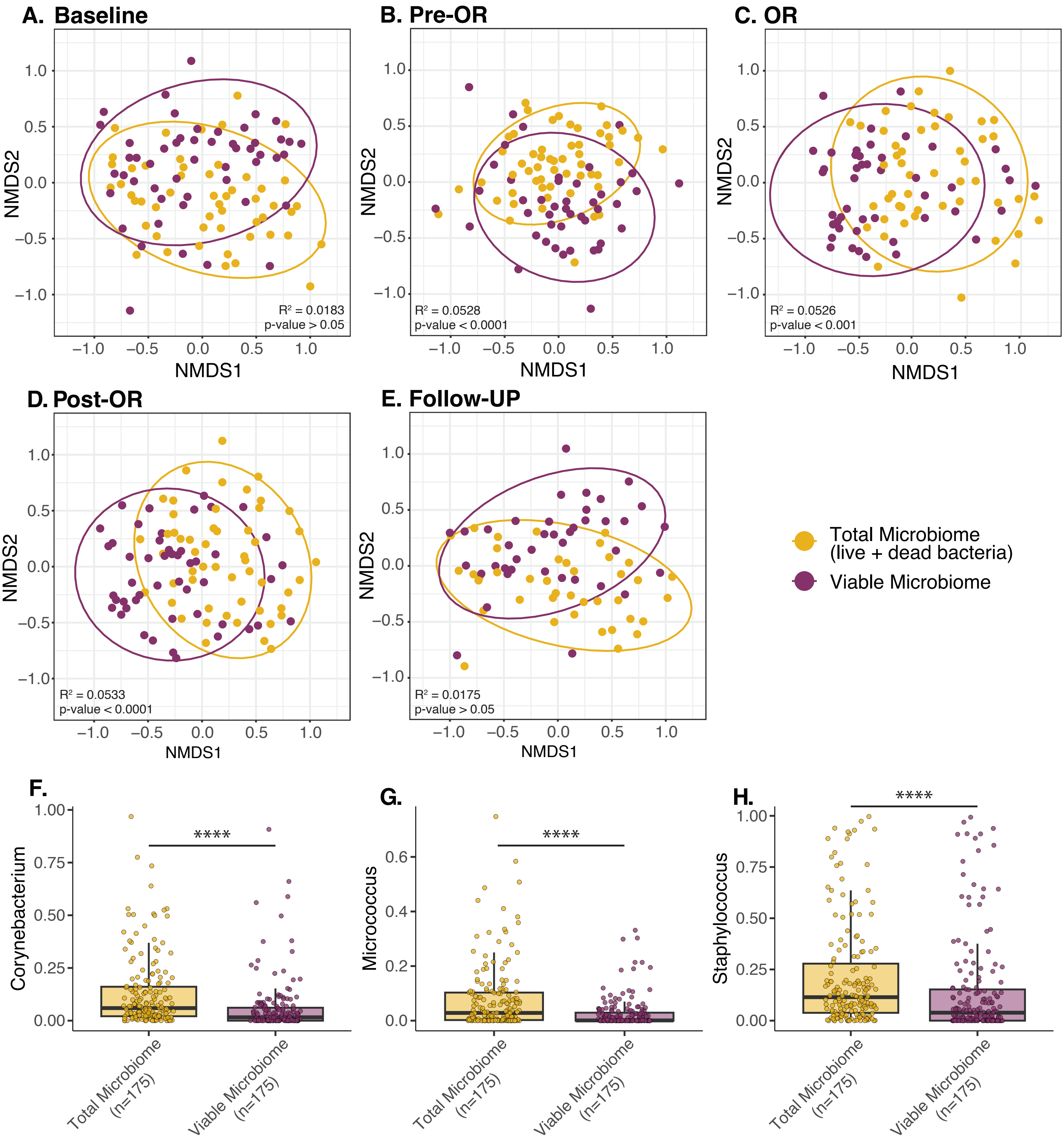


**Supplemental Figure 2: Viable and total microbial community compositions differ on the day of surgery. A-E.** Non-metric Multidimensional Scaling (NMDS) ordination of the Bray-Curtis beta-diversity at each timepoint. PERMANOVAs with 9999 permutations were utilized to evaluate the differences between the viable (PMAxx treated) and total (not treated) sample community compositions. Details can be found in **supplemental table 4. F-H.** MAASLIN2 was used to determine differences in the relative abundance of individual taxa between viable and total communities from samples collected on the day of surgery (Pre-OR, OR, and Post-OR timepoints combined). **** indicates FDR q-value with Benjamini-Hochberg correction < 0.0001.

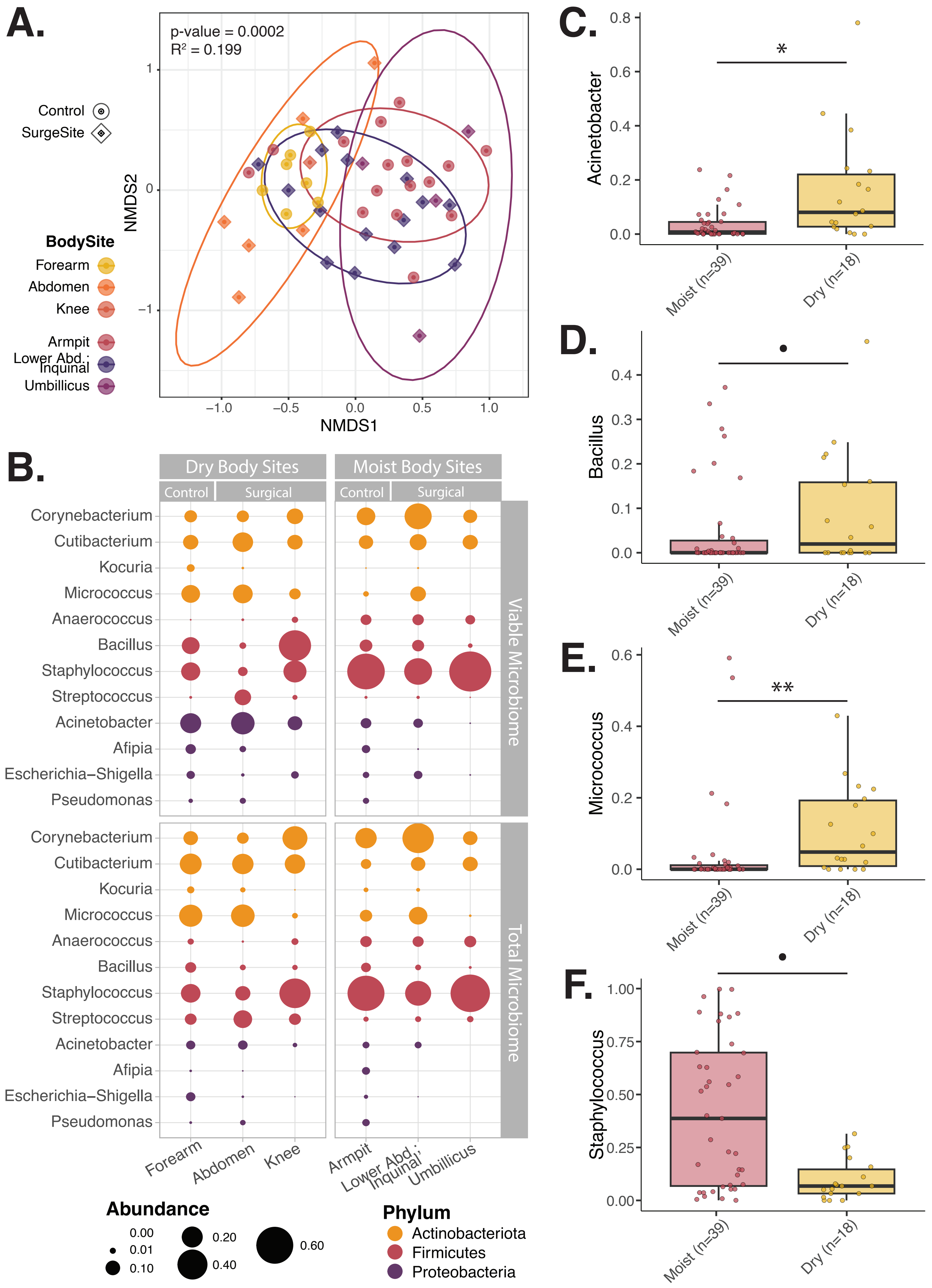
A.
p-value = 0.0002
R² = 0.199
Control
SurgeSite
BodySite
Forearm
Abdomen
Knee
Armpit
Lower Abd.; Inquinal
Umbillicus
NMDS1
NMDS2
B.
Dry Body Sites
Moist Body Sites
Control
Surgical
Corynebacterium
Cutibacterium
Kocuria
Micrococcus
Anaerococcus
Bacillus
Staphylococcus
Streptococcus
Acinetobacter
Afipia
Escherichia–Shigella
Pseudomonas
Viable Microbiome
Total Microbiome
Forearm
Abdomen
Knee
Armpit
Lower Abd.; Inquinal
Umbillicus
Abundance
0.00
0.01
0.10
0.20
0.40
0.60
Phylum
Actinobacteriota
Firmicutes
Proteobacteria
C.
Acinetobacter
D.
Bacillus
E.
Micrococcus
F.
Staphylococcus
Moist (n=39)
Dry (n=18)

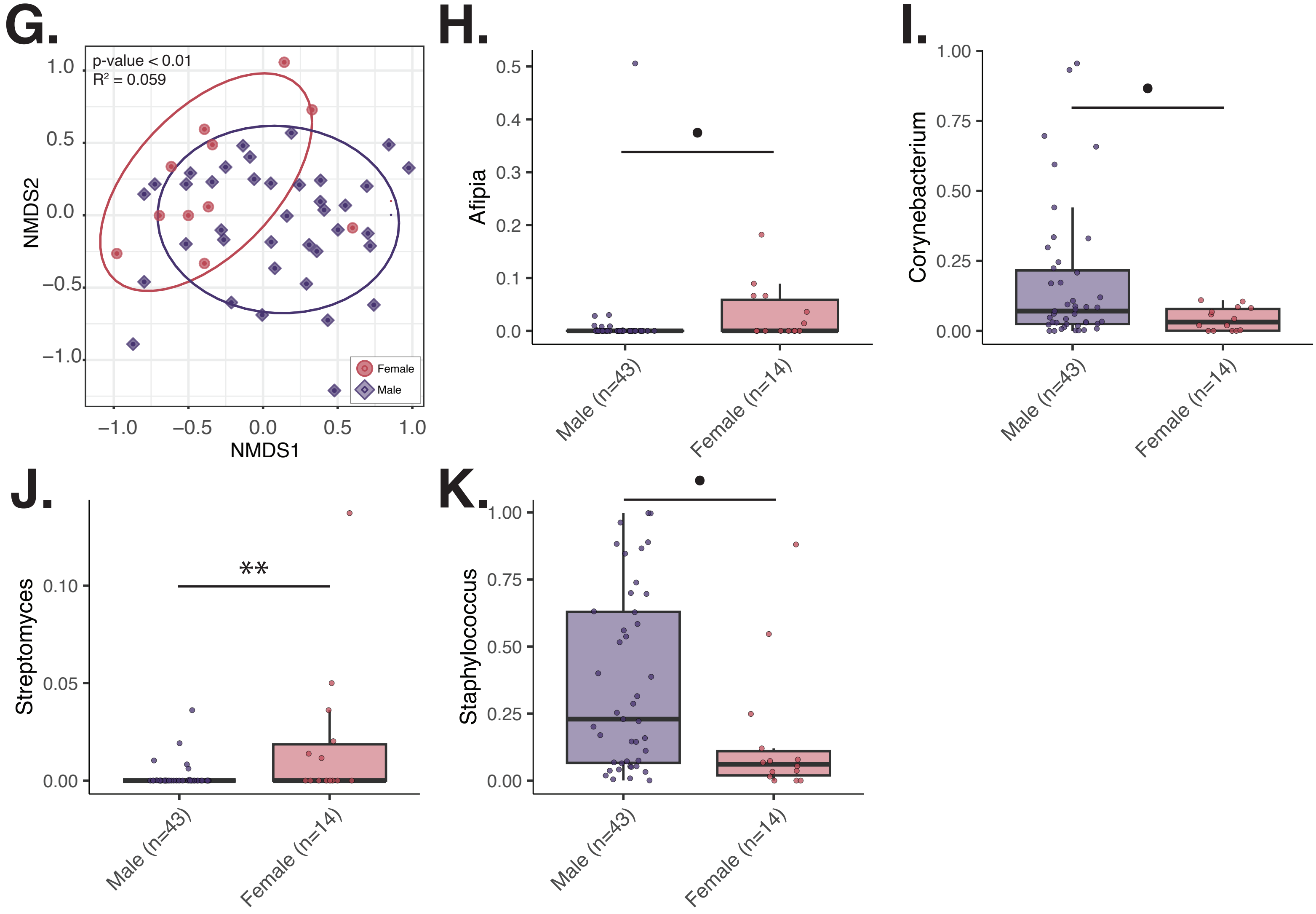


**Supplemental Figure 3: Microbial community composition at baseline is associated with body site of sample collection and subject gender. A.** NMDS ordination of Bray-Curtis beta-diversity of viable microbial communities at subjects' baseline clinic visit highlighting the association between the microbiome composition and body site of sample collection (univariate PERMANOVA with 9999 permutations). **B.** plot of average relative abundance of key genera within viable (top row) and total (bottom row) microbial communities across body sites samples collected at the baseline timepoint. Size of the dot indicates the average genera relative abundance. **C-F.** MAASLIN2 was used to determine differences in the relative abundance of individual taxa within viable communities of samples collected at moist body sites (armpit, lower abdomen to inguinal, and umbilicus combined) compared to those from dry body sites (forearm, central abdomen, and knee combined). Both control and surgical site samples were included in this analysis. In these calculations both subject and gender were incorporated as random effects. Only taxa with significantly different relative abundance ($p$-value $< 0.05$) between the groups are shown. FDR q-values with Benjamini-Hochberg correction indicated. **G.** Bray-Curtis beta diversity NMDS ordination highlighting the association of subject gender with baseline viable microbial community composition (univariate PERMANOVA with 9999 permutations). **H-K.** Differences in the relative abundance of individual taxa within the viable baseline microbiome of male and female subjects were assessed via MAASLIN2. Both control and surgical site samples were included in this analysis and subject and body site of sample collection were incorporated as random effects into these calculations. Only taxa with significantly different relative abundance ($p$-value $< 0.05$) between the groups are shown. FDR q-values with Benjamini-Hochberg correction indicated.

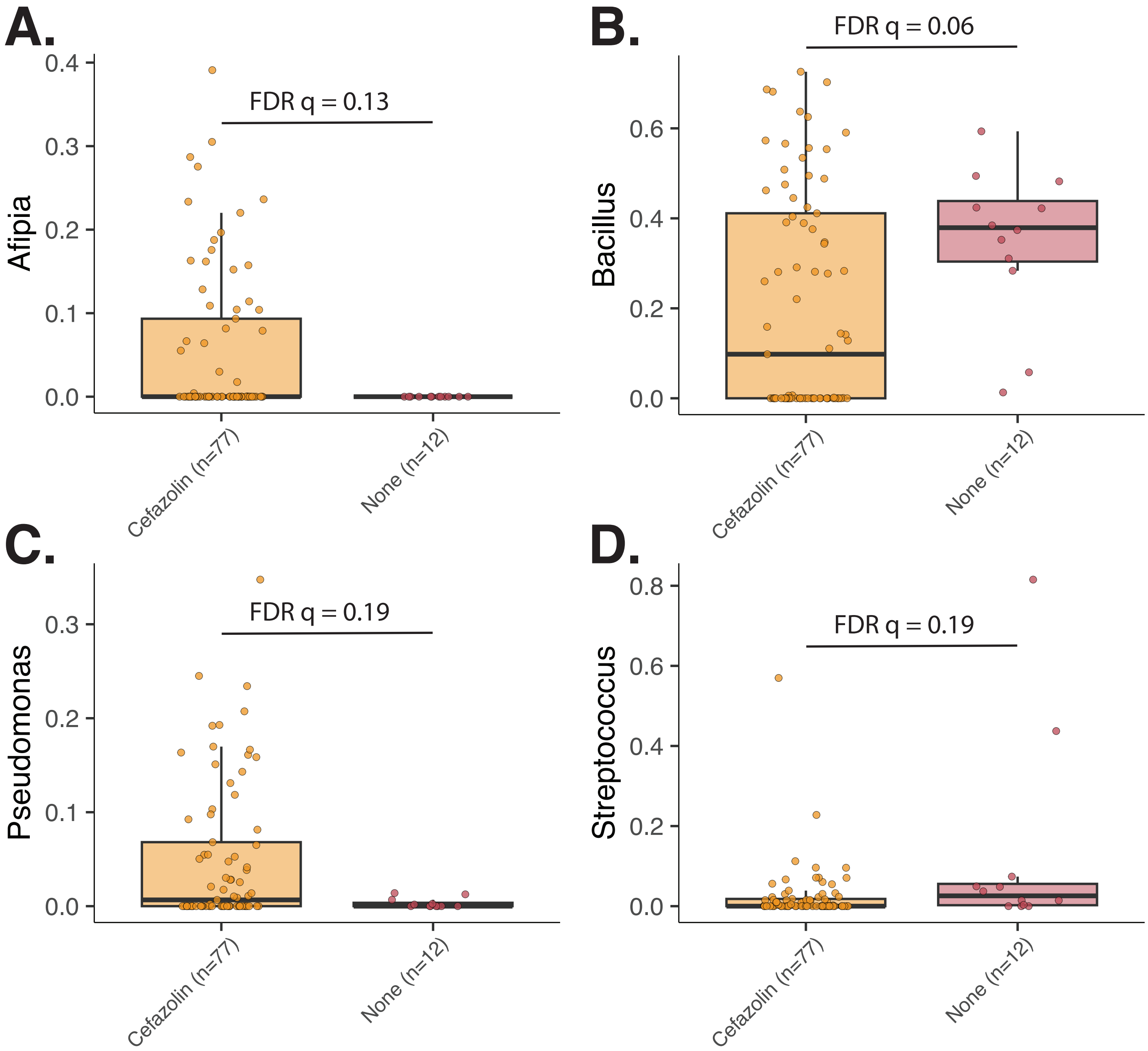


**Supplemental Figure 4: Antibiotic prophylaxis with cefazolin is associated with reduced relative abundance of Bacillus and Streptococcus and increased Afipia and Pseudomonas at the surgical site on the day of surgery. A-D.** Viable microbiome samples from the surgical site on the day of surgery, the samples collected at the pre-OR, OR, and post-OR timepoints, were assessed collectively. Differences in taxa relative abundance were evaluated with MAASLIN2 incorporating the individual subject, gender, and body site of sample collection as random-effects. Only taxa with significantly different relative abundance ($p$-value $< 0.05$) between the groups are shown. FDR q-values with Benjamini-Hochberg correction indicated.

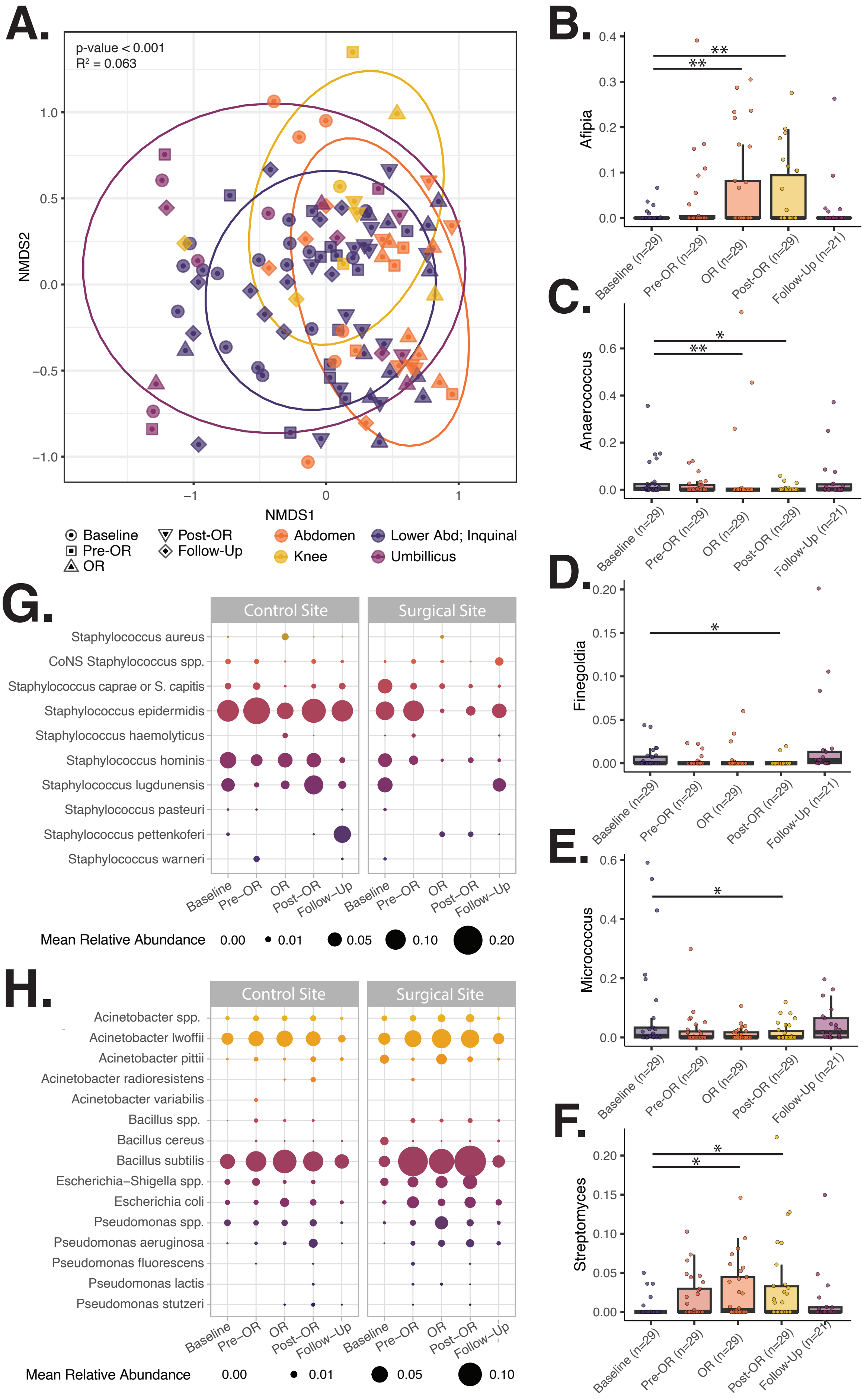


**Supplemental Figure 5: Exposure to Chlorhexidine Gluconate is associated with changes in viable microbial community composition at the surgical site on the day of surgery.** Continuation of **Figure 4. A.** Bray-Curtis beta-diversity NMDS ordination highlighting the association between viable microbial community composition and the body location for the surgical site (univariate PERMANOVA with 9999 permutations). Details can be found in **supplemental table 7. B-F.** Relative abundance plots for taxa significantly more or less abundant in the viable microbiome at the surgery site on the day of surgery, after exposure to CHG, compared to the baseline sample collection. Differential abundance of taxa at each later timepoint compared to the baseline timepoint were assessed via MAASLIN2 and evaluations were made accounting for the individual subject, subject gender, body site of sample collection, and antibiotic prophylaxis as random effects. **Supplemental table 8** contains further details for these results. Note; one subject underwent simultaneous umbilical and inguinal hernia repair. Both sites were sampled at all timepoints. Thus the n = 29 at the baseline through Post-OR timepoints and n=21 at follow-up, which is one more than the total number of subjects who underwent surgery and came for in-person follow-up, n = 28 and = 20 respectively. **G.** ASVs from the Staphylococcus genus were aligned against the BLAST database to obtain probable species assignment. Plot of mean relative abundance of Staphylococcus species within viable microbial communities over time. Apart from S. aureus, all species identified are coagulase negative Staphylococci (CoNS). “CoNS Staphylococcus spp.” indicates ASVs that aligned well to several CoNS species. **H.** ASVs from the Acinetobacter, Bacillus, Escherichia-shigella, and Pseudomonas genera were aligned against the BLAST database to obtain probable species assignment. Plot displays mean relative abundance of these species within viable microbial communities over time.

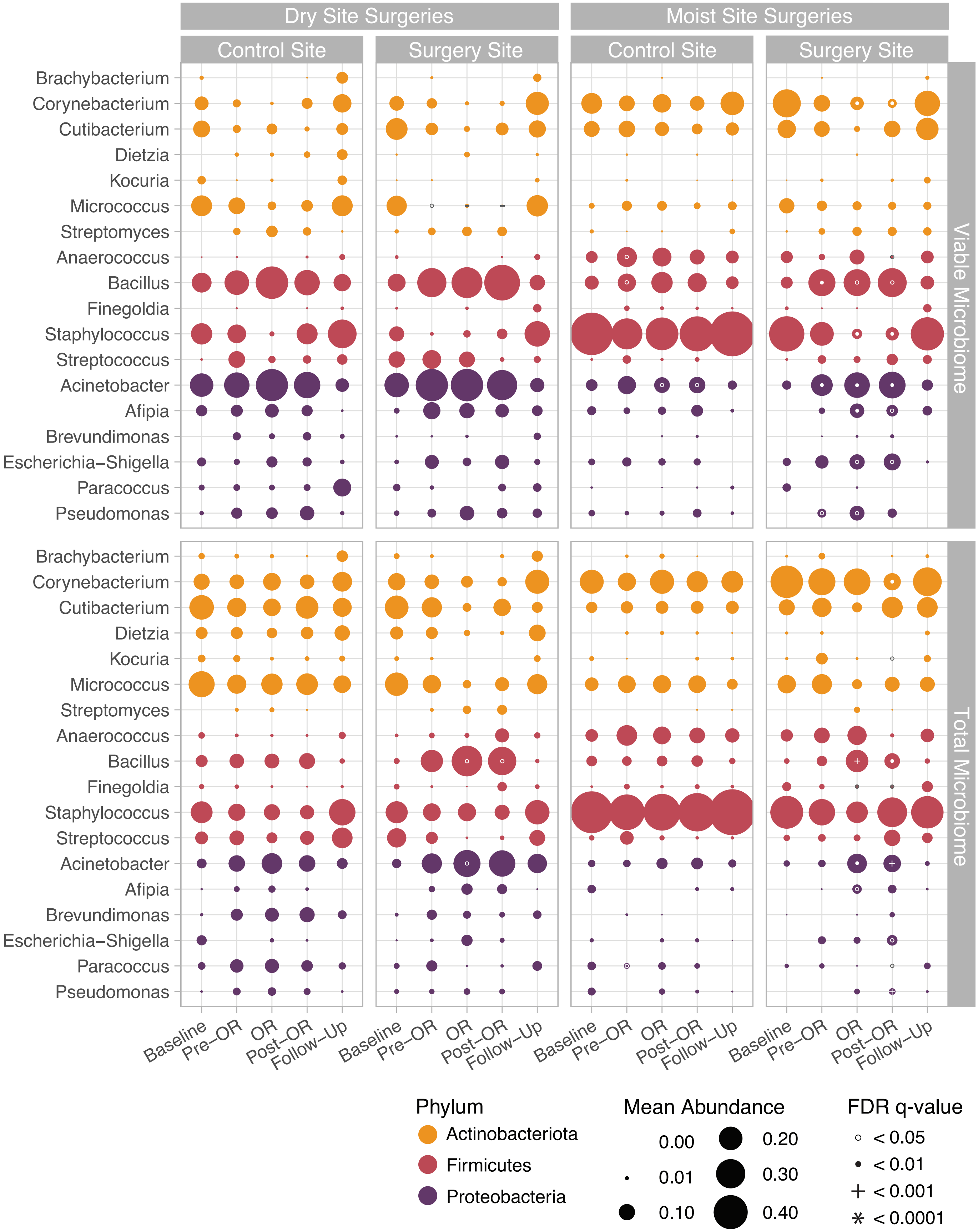


**Supplemental Figure 6: Change in average relative abundance of key taxa over time at surgical and control sites collected from most and dry body sites.** Companion to **Figure 4C** displaying the average relative abundance of each taxa over time in the viable and total microbial communities at moist and dry surgical sites and control sites respectively. Mean taxa abundance is indicated by the size of the point. Differential relative abundance of taxa at later timepoints versus at baseline was evaluated via MAASLIN2. All MAASLIN evaluations were made accounting for subject, body site of sample collection, gender, and antibiotic prophylaxis as random effects. White, or in a few cases grey, circles, filled-in dots, plus sign, and asterixis indicate the degree of significance. Further details for the MAASLIN2 results located in **Supplemental Table 8.**

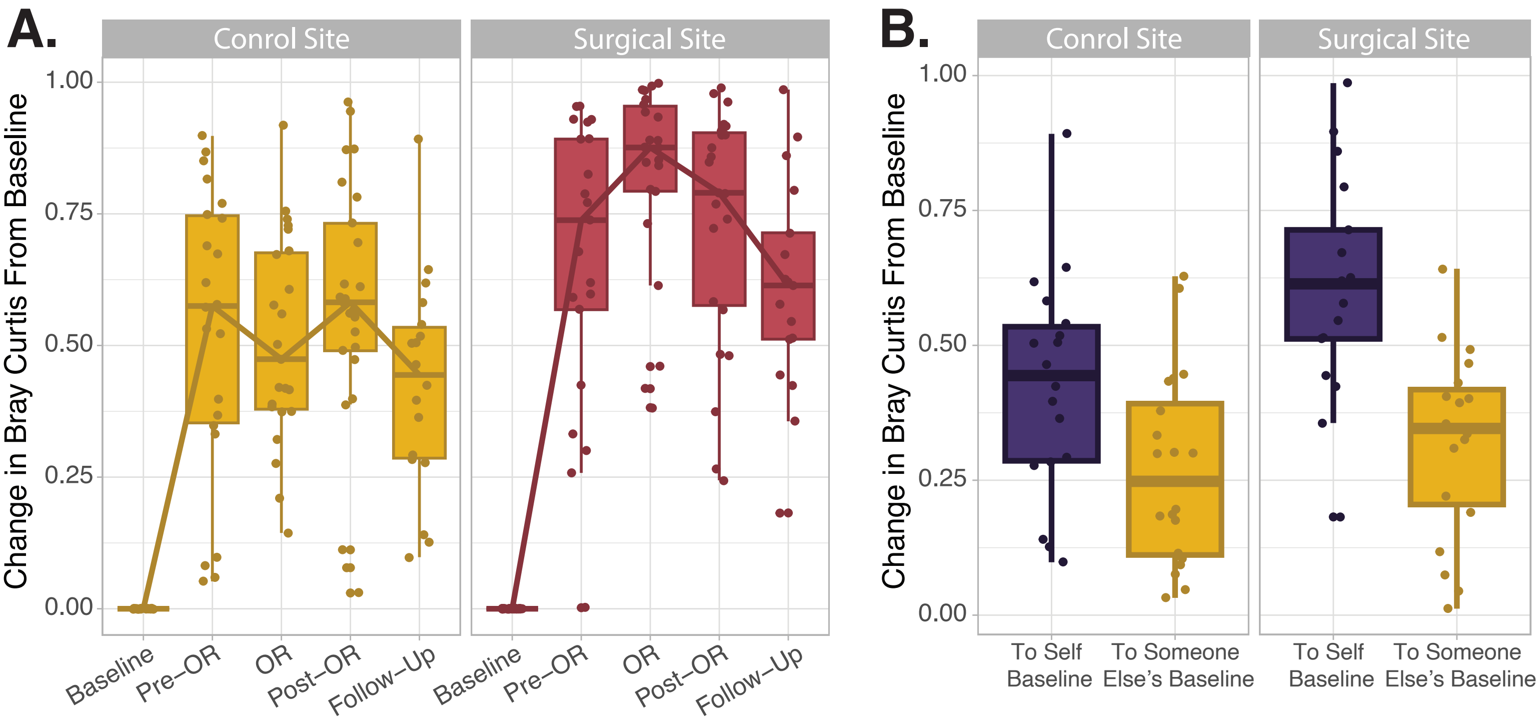


**Supplemental Figure 7: Change microbial community composition from baseline. A.** Similarity of each subject's surgical or control site microbiome at each timepoint was compared to their respective baseline community composition via the Bray-Curtis beta diversity metric. **B.** Bray Curtis beta diversity metric was also used to evaluate the similarity of each subject's microbiome composition at follow-up to both their own baseline microbiome composition and the most similar baseline microbial community composition (smallest Bray-Curtis distance) of another subject. Differences between the bray-curtis distance between a subject's follow-up to their own baseline versus someone elses baseline microbiome were evaluated with the Wilcoxon matched pairs signed rank tests. ** indicates p-value < 0.01; **** indicates p-value < 0.0001.

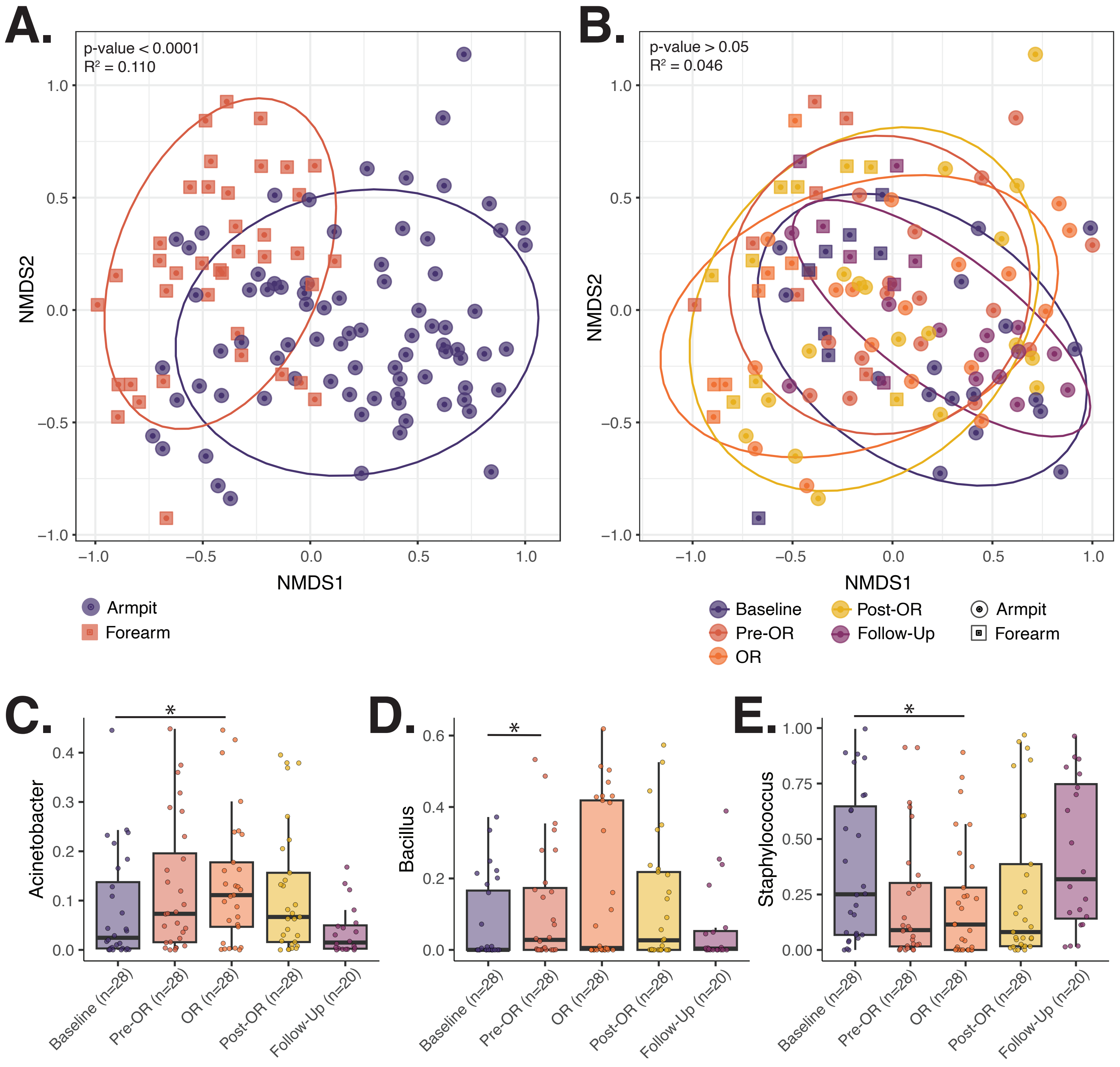


**Supplemental Figure 8: Viable communities at the control sites are more associated with the body site of sample collection.** Companion figure to **figure 4** and **supplemental figure 5**. **A.** Bray-Curtis beta diversity NMDS ordination illustrating the strong association of body site of control samples with viable microbial community composition. **B**. Bray-Curtis beta-diversity ordination highlighting the lack of association between viable microbial community composition and timepoint of control site sample collection. Associations of microbial community composition with various features were evaluated via univariate PERMANOVAs with 9999 permutations. Details can be found in **supplemental table 7. C-E.** Relative abundance of Acinetobacter (C), Bacillus (D), and Staphylococcus (E) in viable communities at both moist (armpit) and dry (forearm) control sites over time. Differential abundance of taxa at each timepoint compared to the baseline timepoint were assessed via MAASLIN2. All MAASLIN2 evaluations were made accounting for individual subjects, gender, body site of sample collection and antibiotic prophylaxis as random effects.

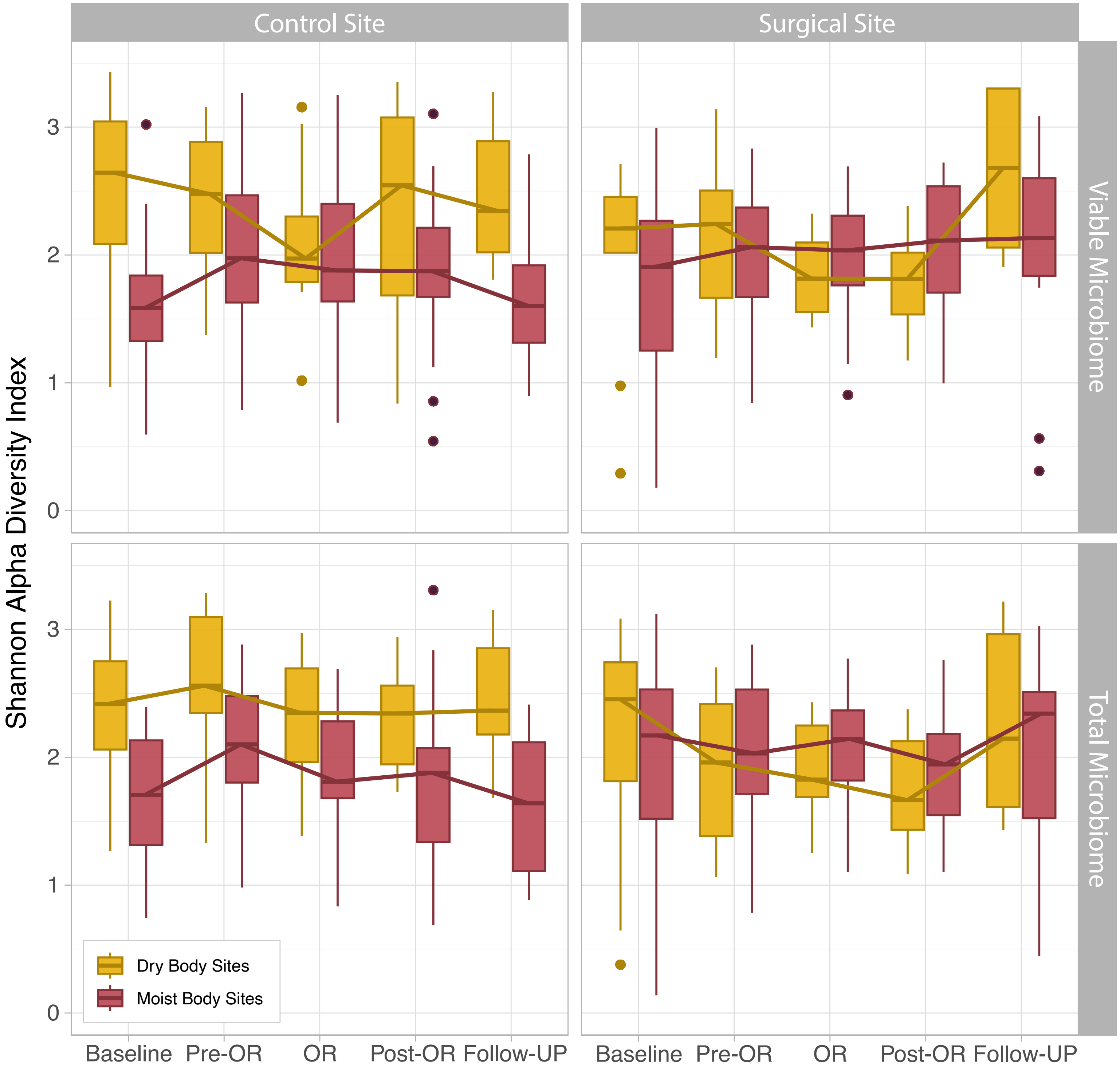


**Supplemental Figure 9: No change in Shannon alpha diversity following CHG application.** Shannon alpha diversity metric was used to measure the microbial diversity within each sample. Boxplots represent the Shannon index distribution (median interquartile range) of viable and total microbial communities at the surgical and control sites. Samples are grouped by if they were collected at a moist or dry body site.
